# Regulators of health and lifespan extension in genetically diverse mice on dietary restriction

**DOI:** 10.1101/2023.11.28.568901

**Authors:** Andrea Di Francesco, Andrew G. Deighan, Lev Litichevskiy, Zhenghao Chen, Alison Luciano, Laura Robinson, Gaven Garland, Hannah Donato, Will Schott, Kevin M. Wright, Anil Raj, G.V. Prateek, Martin Mullis, Warren Hill, Mark Zeidel, Luanne Peters, Fiona Harding, David Botstein, Ron Korstanje, Christoph A. Thaiss, Adam Freund, Gary A. Churchill

## Abstract

Caloric restriction (CR) delays aging and extends healthy lifespan in multiple species. Alternative forms of dietary restriction (DR) such as intermittent fasting (IF) have drawn significant interest as a more sustainable regimen, but the landscape of longevity-promoting dietary interventions remains largely unexplored. Identifying the most robust, efficacious, and experimentally tractable modes of DR is key to better understanding and implementing effective longevity interventions for human healthspan. To that end, we have performed an extensive assessment of DR interventions, investigating the effects of graded levels of CR (20% and 40%) and IF (1 day and 2 days of fasting per week) on the health and survival of 960 genetically diverse female mice. All interventions extended lifespan, although only CR significantly reduced the mortality doubling time. Notably, IF did not extend lifespan in mice with high pre-intervention bodyweight. We carried out extensive phenotyping to determine the health effects of long-term DR and to better understand the mechanisms driving within-diet heterogeneity in lifespan. The top within-diet predictor of lifespan was the ability of mice to maintain bodyweight through periods of handling, an indicator of stress resilience. Additional predictors of long lifespan include specific changes in immune cells, red blood cell distribution width (RDW), and retention of adiposity in late life. We found that lifespan is heritable (h^2^ = 0.24), and that genetic background has a larger influence on lifespan than dietary interventions. We identified a significant association for lifespan and RDW on chromosome 18 that explained 4.3% of the diet-adjusted variation in lifespan. Diet-induced changes on metabolic traits, although beneficial, were relatively poor predictors of lifespan, arguing against the long-standing notion that DR works by counteracting the negative effects of obesity. These findings indicate that improving health and extending lifespan are not synonymous and that metabolic parameters may be inappropriate endpoints for evaluating aging interventions in preclinical models and clinical trials.

## Introduction

Caloric restriction (CR) delays the onset of age-related diseases and extends lifespan in multiple species^1^. In humans, compliance with CR is challenging, and interest has shifted to more permissive forms of dietary restriction (DR), such as time-restricted feeding and intermittent fasting (IF) that have proven to be effective in promoting organismal health^2–5^. In mice, regular periods of fasting can convey significant benefits without reduction in overall energy intake^6^. Mice on CR also experience prolonged periods of daily fasting, and the health benefits of CR can be optimized by feeding at a specific time of the day, suggesting that both caloric intake and circadian feeding patterns contribute to physiological response and lifespan extension^5,7,8^. Despite the importance of these observations, limited information is available regarding the differences between CR and IF in relation to aging and longevity^9^.

Responses to DR vary across individuals, and the mechanisms underlying this variability remain largely unknown. Studies in mice and non-human primates have shown that the effects of DR are influenced by individual characteristics including sex, body size and composition, and genetic background^8,10–17^. Human clinical studies testing the effects of DR on health parameters have largely focused on changes in body weight, adiposity, energy metabolism, and cardiometabolic risk factors^18–23^. There has been less exploration of the long-term effects of DR because studies in humans are limited by their small sample size and short duration. Additionally, the safety and efficacy of DR may depend on factors such as age and health condition^24^. There is a lack of knowledge regarding physiological markers that can be used to predict how individuals will respond to DR. Discovery of such predictors could help tailor DR to individual needs, serve as tools in the longitudinal evaluation of intervention success, and elucidate the biological processes that mediate their effects on lifespan.

In this study, we investigated the effects of CR and IF on the health and lifespan of female Diversity Outbred (DO) mice^25^. These genetically diverse mice present a wide range of physiological characteristics, including variation in body composition and other metabolic traits,^26^ which could serve as predictive markers of individual response to DR. Findings from genetically diverse populations are more likely to generalize across species, reveal heterogeneity in response to DR, and enable the identification of the genetic basis of trait variation. Recent studies incorporating a different genetically diverse mouse population have revealed substantial variation in normative aging^27^. Our study of Dietary Restriction in Diversity Outbred mice (DRiDO) longitudinally collected a broad range of health-related outcomes to reveal beneficial and detrimental effects of DR, and identifies physiological and genetic factors that predict individual response to DR.

### Study design

We examined the effects of graded levels of CR and IF on 960 female DO mice that were randomly assigned to one of five diets: *ad libitum* feeding (AL), fasting one day per week (1D) or two consecutive days per week (2D), and caloric restriction at 20% (20) or 40% (40) of baseline *ad libitum* food intake (**Fig. 1a**). At 6 months of age, we initiated DR on 937 surviving mice and maintained them on DR for the duration of their natural lifespan.

**Fig. 1.**
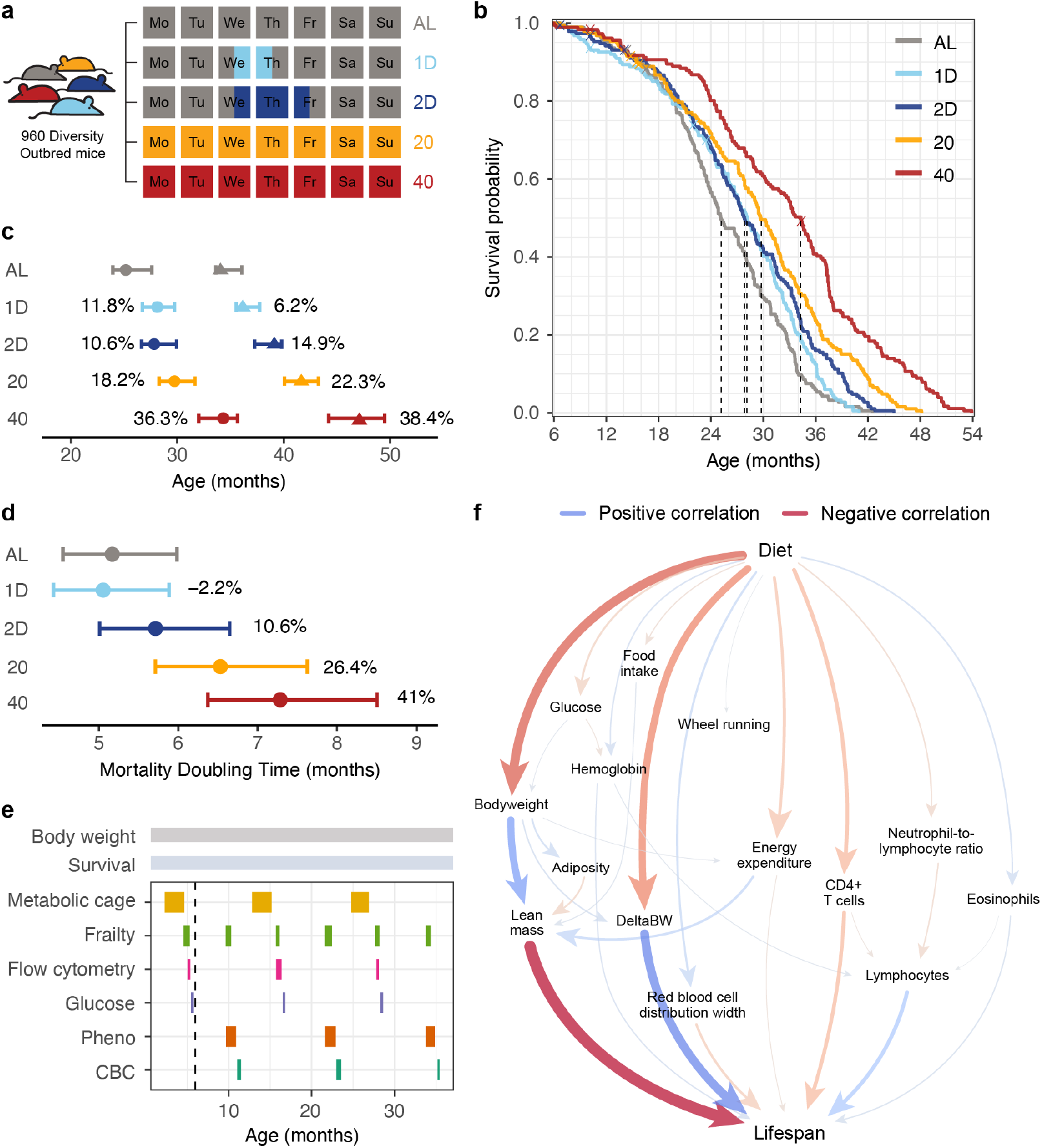
Dietary restriction extends lifespan in DO mice. **a**, Overview of feeding schedules. **b**, Kaplan-Meier survival curves by diet group. Dashed lines indicate median survival. **c**, Median and maximum lifespan by diet group with 95% confidence intervals and percent change relative to AL. **d**, Mortality doubling times estimated from a Gompertz log-linear hazard model with 95% confidence interval and percent change relative to AL indicated. **e**, Phenotyping schedule. Phenotypes (Pheno) include echocardiogram, body composition, rotarod, acoustic startle, and wheel running. A blood draw for complete blood count (CBC) was taken at the end of the phenotyping period. **f**, Waterfall plot shows the highest scoring paths from Diet to Lifespan in the partial correlation network. Size of the arrow is proportional to absolute partial correlation and color indicates the sign of the partial correlation.

We used an additional 160 DO mice to establish the distinct daily and weekly feeding patterns induced by this dietary protocol (**Extended Data Fig. 1a-c**). IF mice had their food removed at 3pm on Wednesday and replenished after 24 (1D) or 48 (2D) hours. The IF mice experienced 6 to 10% bodyweight loss while fasting (**Extended Data Fig. 1d**). IF mice displayed compensatory feeding following their fasting period such that 1D mice consumed a similar amount of food as AL mice, while 2D mice consumed approximately 12% less food over the course of one week. CR mice were fed a measured amount of food daily at 3pm. On Friday afternoon, CR mice received a triple amount of daily food allotment that was typically consumed by 3pm on Saturday (40% CR) or 3pm on Sunday (20% CR), resulting in weekly fasting periods like those experienced by the 2D and 1D mice, respectively. In contrast to the IF mice, the CR mice showed no change in bodyweight over the weekend despite experiencing a period of fasting. These dietary interventions had substantial effects on metabolism, energy expenditure, and activity of mice (**Extended Data Fig. 2, 3**).

### DR extends lifespan in female DO mice

DR extended the lifespan of female DO mice (log rank p < 2.2e-16), with response proportional to the degree of restriction or length of fasting (40% > 20% > 2D > 1D > AL; **Fig. 1b; Supplementary Table 1**). Divergence among the lifespan curves was not apparent until ∼18 months of age, suggesting that intervention effects on survival accumulate gradually or have a delayed onset. DR affected both median and maximum lifespan (50% and 90% mortality, respectively^28^) (**Fig. 1c**; **Supplementary Table 2**). 40% CR mice achieved a median lifespan ∼9 months (36.3%) greater than mice in the AL group. We estimated the mortality doubling time by fitting Gompertzian lifespan models^29^ to the post-DR survival data (**Fig. 1d, Supplementary Table 3**) and observed a significant decrease in the rate of aging for CR mice but not for IF mice compared to AL.

### Regulators of DR-mediated lifespan extension

Despite the profound effects of DR, lifespan was highly variable within intervention groups (**Extended Data Fig. 4a**). To identify possible regulators of lifespan variability, we carried out longitudinal phenotyping across multiple domains of mouse physiology (**Fig. 1e**). We obtained weekly body weights; assessed frailty index, grip strength, and body temperature every 6 months; and carried out yearly assessments including metabolic cage analysis, body composition, echocardiogram, wheel running, rotarod, acoustic startle, bladder function, fasting glucose, immune cell profiling, and whole blood analysis. We evaluated a total of 194 traits, resulting in up to 689 single timepoint measurements per mouse. We also analyzed the gastrointestinal microbiome by collecting fecal samples every 6 months; a complete analysis of the microbiome data is available in a companion paper (see Litichevskiy et al.)

We performed three distinct analyses (see Methods). First, longitudinal analysis was used to assess how traits change with age and diet (**Extended Data Fig. 4b, Supplementary Table 4**). Second, correlation analysis was used to identify traits that predict lifespan (**Extended Data Fig. 4c-e, Supplementary Table 5**). Third, multivariate network analysis was used to identify traits that potentially mediate the effects of DR on lifespan. For the network analysis, we first fit a partial correlation network (sparse undirected Gaussian graphical model^30^) to all 194 traits (**Extended Data Fig. 5a; Supplementary Table 6**) and then fit a new partial correlation network to a reduced set of 21 exemplar traits. This analysis revealed that the top scoring pathways for potential mediation between diet and lifespan fall into two broad categories: (1) bodyweight and body composition, and (2) immune cell composition and hematologic traits (**Fig. 1f, Extended Data Fig. 5b**). Notably, the effects of DR are not universally life-extending – detrimental contributions are associated with a loss of ability to maintain bodyweight and with specific changes in immune and hematologic traits, including the coefficient of variation for red blood cell distribution width (RDW), as discussed below.

Below, we look at traits that predict lifespan and individual response to DR. We first examine bodyweight and body composition, then we turn to health and metabolic traits, and finally immune and hematological traits. Lastly, we examine the influence of genetics.

### Bodyweight and lifespan prediction

DR has a profound effect on body weight (**Fig. 2a**). We observed substantial variation in bodyweight trajectories among individual mice (**Extended Data Fig. 6a**). Changes in growth trajectories were apparent from the onset of DR. Notably, 1D mice lost weight despite having a similar caloric intake as the AL mice. The 40% CR group was distinguished by a rapid loss of bodyweight, losing an average of 21.8% of their 6-month bodyweight by 20 months of age. In contrast, AL mice gained 28.5% bodyweight over the same period. Weight loss in the 40% CR mice persisted throughout life, suggesting that most of these animals never achieved energy balance. To compare mice at similar life stages and remove survivorship bias, we rescaled age from chronological time to proportion of life lived (PLL = age at test / lifespan), a surrogate for biological age. Replotting the bodyweight trajectories as a function of PLL (**Fig. 2b**) revealed that for groups other than 40% CR, weight gain continues through midlife, stabilizes between 0.50 to 0.75 PLL, and declines rapidly near the end of life (beyond 0.90 PLL).

**Fig. 2.**
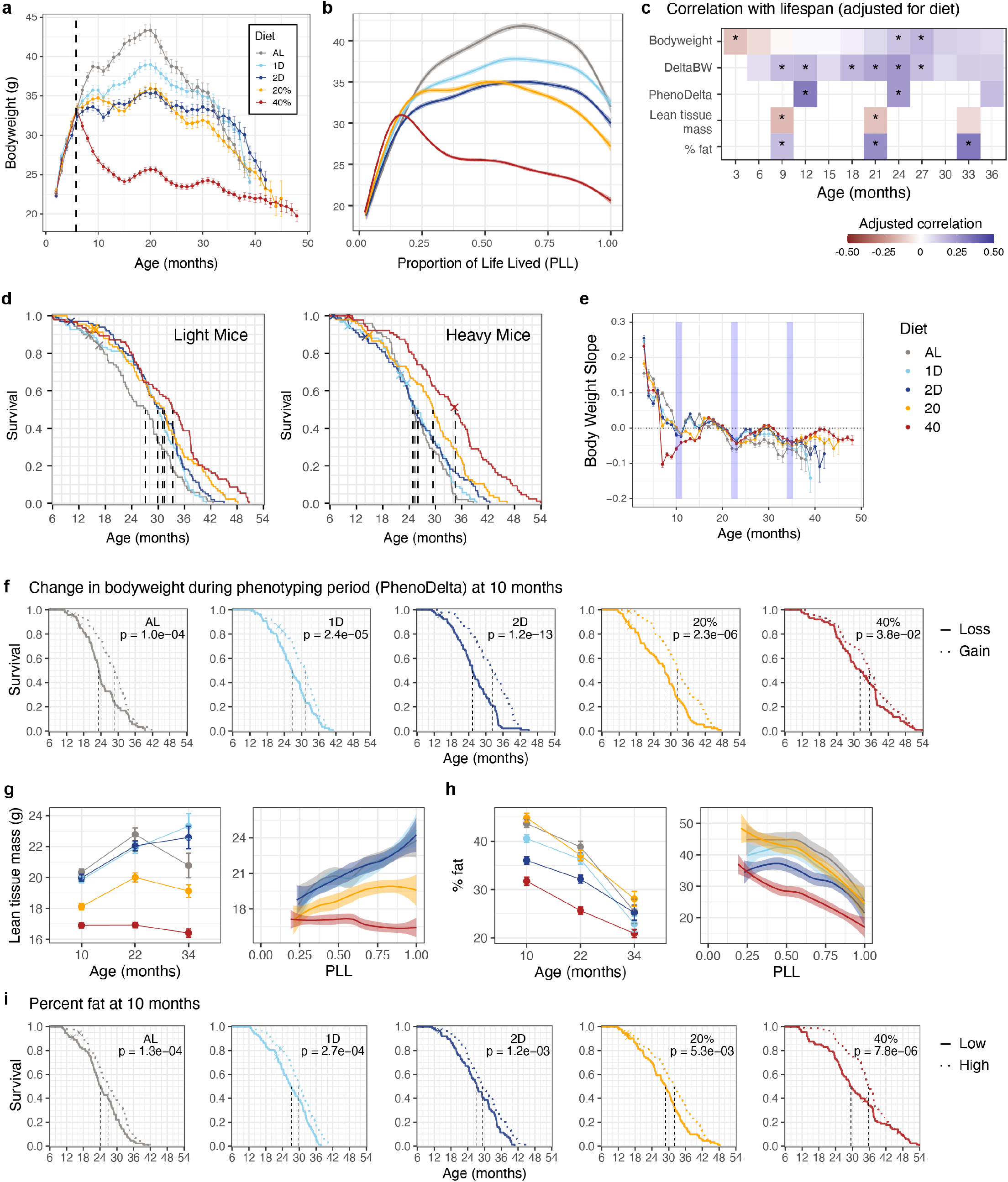
Post-DR bodyweight and adiposity predict lifespan. **a,** DR alters the lifetime trajectory of bodyweight. Individual-mouse weekly bodyweight data were loess smoothed and monthly means (± 1SE) were plotted for each diet group. **b,** Bodyweight (g) trajectories are plotted against proportion of life lived (PLL) as loess smoothed curves with 95% confidence bands. **c,** Heatmap showing Pearson correlation with lifespan for bodyweight, change in bodyweight across three-month intervals (DeltaBW), change in bodyweight across one-month phenotyping windows (PhenoDelta), lean tissue mass, and adiposity (% fat). Asterisks indicate significant association between the trait and lifespan (FDR < 0.01). Correlations were computed from rank normal scores transformed data and are adjusted for diet. **d**, Kaplan-Meier survival curves by diet group (color) are shown for mice below and above median 6-month bodyweight. Dashed lines indicate median survival time per group. **e**, Monthly slopes (log of proportional change in bodyweight per month) of individual mouse growth trajectories were averaged within diet groups (mean ± 1SE) and plotted against time. Values above zero indicate that, on average, mice are gaining weight, and values below zero indicate average weight loss. Blue vertical bands indicate the time-windows in which phenotype testing was carried out. **f,** Kaplan-Meier curves for each diet group compare the survival of mice by diet-specific change in bodyweight across the 10-month phenotyping period (PhenoDelta 10). **g, h,** Lean tissue mass (g) and adiposity (h) are plotted against age (mean ± 2SE) and against PLL (loess smooth with 95% confidence band). **i**, Kaplan-Meier curves within each diet group compare survival of mice stratified by diet-specific median adiposity at 10 months of age (percent fat at 10 months). Panels f and i show as inset the uncorrected p-value from within-diet linear regression of survival time on continuous trait values (transformed to rank normal scores).

Early body weight has been shown to predict lifespan in mice and other species^31–34^. Consistently, we observed a negative correlation between lifespan and bodyweight in early life (**Fig. 2c**, top row). This association decreases with age and becomes positive beyond two years of age. We asked if early bodyweight might also modify response to DR. Kaplan-Meier analysis stratified by median 6-month bodyweight showed that CR extended lifespan regardless of pre-DR bodyweight and there was no significant difference in lifespan extension between lighter and heavier mice (**Fig. 2d, Extended Data Fig. 6b, Supplementary Table 5**). In contrast, we observed that lighter mice benefited from IF, but heavier mice did not respond to IF and had lifespans comparable to AL mice.

We asked whether individual lifespan extension could be explained by weight loss in the post-DR growth phase between 6 and 20 months of age^35^. On the contrary, we found that within diet groups, mice that retained more weight had longer lifespans (overall p = 1.77e-10) (**Extended Data Fig. 6c**). We also looked at change in bodyweight at three-month intervals (DeltaBW) and found that weight loss at any age was associated with reduced lifespan (**Fig. 2c**, second row).

During the one-month long phenotyping periods at 10, 22, and 34 months of age, mice were subject to intensive characterization and experienced stress due to handling and housing changes. Most mice experienced short-term weight loss followed by recovery in the following weeks (**Fig. 2e**). Resilience to change in bodyweight from 10 to 11 months of age (PhenoDelta 10) was the top predictor of lifespan across all traits examined in this study (p < 2.2e-16; **Fig. 2c**, third row; **Fig. 2f**). Weight loss during the 22-month-old phenotyping period was also associated with reduced lifespan (**Extended Data Fig. 6d**).

To better understand whether changes in body composition contributed to these effects on lifespan, we analyzed the body composition of mice at 10, 22, and 34 months of age. We deconvolved total tissue mass into lean tissue mass and adiposity (percent fat). The genetically diverse DO mice displayed wide variation in bodyweight and composition, with adiposity ranging from less than 10% to over 60% across all diet groups and ages (**Extended Data Fig. 7a**). For AL and IF mice, lean tissue mass increased throughout life along identical trajectories on the PLL scale (**Fig. 2g**). For CR mice, lean mass was reduced compared to AL mice and remained constant (20%) or declined (40%) with age. All groups of mice lost fat mass with age, particularly in the latter half of life (>0.5 PLL) (**Fig. 2h**). While 40% CR mice had the lowest average adiposity, 20% CR mice had adiposity levels comparable to AL mice, and individual mice with the highest adiposity were found in the 20% CR group.

Previous studies have shown that preserving fat mass may increase survival in response to chronic CR^11,16,36^. We examined the association between lifespan and adiposity at 10 (**Fig. 2i**) and 22 months of age (**Extended Data Fig. 7b**). Higher adiposity was consistently associated with increased lifespan across diets and age, and the association was stronger in later life (10mo: p = 5.28e-15; 22mo: p < 2.2e-16). The most pronounced differences were seen in the 40% mice as early as 10 months of age, and higher adiposity was significantly beneficial across all diet groups by 22 months. Lean tissue mass was negatively correlated with lifespan, more strongly early in life, while adiposity was positively correlated, more prominently in late life (**Fig. 2c**, bottom two rows). The net effect of these two trends provides a plausible explanation for the reversal of bodyweight-lifespan association with age. Collectively, these results highlight the critical role of body weight and composition on the outcome of different DR interventions.

### Impact of DR on health and metabolism

Many of the traits in our phenotyping pipeline were selected to evaluate the health of aging mice. Surprisingly, a relatively modest number of them showed associations with lifespan (**Fig. 3a, b, Extended Data Fig. 4c**). The frailty index^37^ (FI) is an indicator of morbidity that measures age-related health deficits. As expected, the FI score increased with age (**Fig. 3c**). AL mice exhibited the highest number of FI events by chronological age, but on the PLL scale there was no significant difference between diet groups in the rate of accumulation, suggesting that FI is a reliable marker of biological age.

**Fig. 3.**
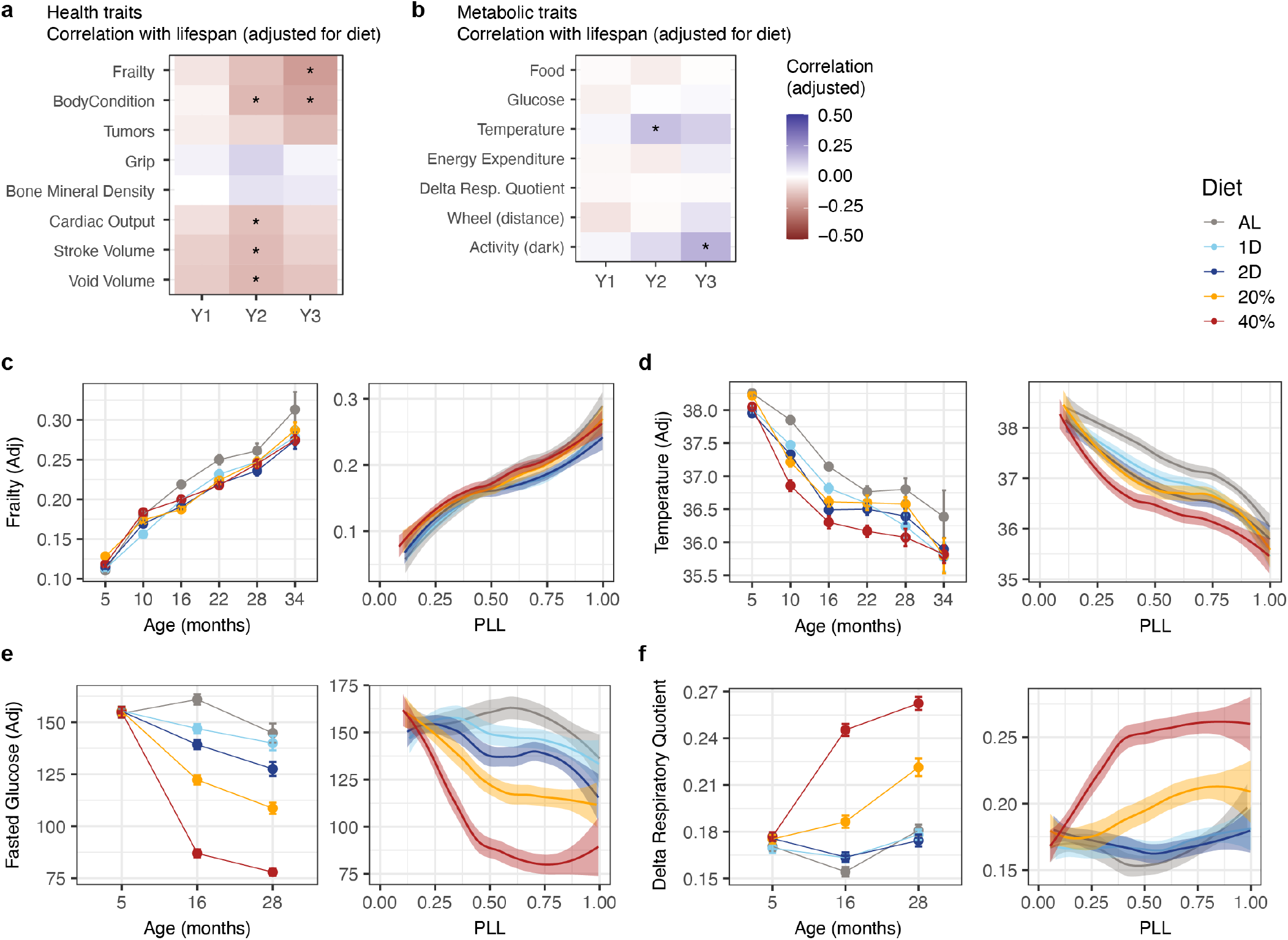
Health and metabolic traits change with age and diet but are poor predictors of lifespan. **a, b**, Heatmaps show the Pearson correlation with lifespan for (a) selected health-related traits and (b) selected metabolic traits (y-axis) at annual testing intervals (x-axis). Asterisks indicate significant association between the trait and lifespan (FDR < 0.01). Correlations were computed from rank normal scores transformed data and adjusted for diet. **c-f**, The frailty index adjusted for batch effects (c), body temperature adjusted for bodyweight (d), fasted glucose levels adjusted for bodyweight (e), and average daily change in respiratory quotient (Delta Respiratory Quotient, f) are plotted against age (mean ± 2SE) and against PLL (loess smooth with 95% confidence band).

Individual FI components revealed diet-specific beneficial effects including reduced incidence of palpable tumors and distended abdomen, as well as detrimental effects on body condition and increased incidence of kyphosis among 40% CR mice (**Extended Data Fig. 8a-d**). We observed declines in bodyweight-adjusted grip strength and bone mineral density with age, but DR had little effect on these traits (**Extended Data Fig. 8e, f**). In addition to FI, the echocardiogram traits stroke volume and cardiac output, and total volume of urine in the voiding assay showed modest associations with lifespan (**Fig. 3a**).

We also observed long-term metabolic responses to DR. Mouse body temperature declined with age, and DR resulted in a further reduction that was maintained throughout life (**Fig. 3d**). Fasting glucose levels peaked at 16 months in AL mice, declined at later age, and were significantly reduced by DR (**Fig. 3e**). The difference in respiratory quotient between day and night (Delta RQ), which mirrors the metabolic transition between fed and fasted states, was more pronounced in mice subjected to 20% and 40% CR compared to both the AL and IF groups (**Fig. 3f**). Energy expenditure (EE) adjusted for body weight declined from pre-intervention levels and plateaued, with lowest levels in 40% followed by 20% and 2D groups (**Extended Data Fig. 8g**). Wheel running activity declined with age across all groups, with the notable exception of 40% CR mice, which maintained elevated activity levels throughout their lifetime, a consequence of persistent food seeking behavior^38,39^ (**Extended Data Fig. 8h**).

Improved glucose homeostasis, lower EE, decreased body temperature, and preservation of metabolic flexibility (Delta RQ) are common physiological adaptations to DR in both rodents and humans that have been suggested as potential mechanisms that mediate the extension of lifespan associated with DR^40,41^. However, we found no significant association between lifespan and fasting glucose, EE, or Delta RQ at any age. Higher body temperature was moderately associated with increased lifespan, as was activity in the dark phase, but overall, measures of metabolic health and fitness displayed only weak or no associations with lifespan (**Fig. 3b**; **Extended Data Fig. 8i-j**). These observations indicate that, unexpectedly, many of the health benefits of DR are not mediators of longevity effects.

### Impacts of DR on immune and erythroid cell composition

In contrast to health and metabolic traits, we found that after adjusting for diet, many immune and hematological traits were correlated with lifespan (**Fig. 1f, 4a, b, Extended Data Fig. 4c, d**). Age-related changes in immune cell subsets in DO mice generally aligned with changes previously described in humans and in common inbred mouse strains^42–45^. Overall, B cells, effector T cells and inflammatory monocytes accumulated with age, while total lymphocytes, mature NK cells and eosinophils declined. DR did not significantly modify the age-related changes in the percent of lymphocytes or the percent of effector T cells (**Fig. 4c, d, Extended Data Fig. 9a, b**). However, 40% CR had a profound effect on the frequency of mature NK cells, eosinophils, circulating B cells, and inflammatory monocytes (**Extended Data Fig. 9c-f**).

**Fig. 4.**
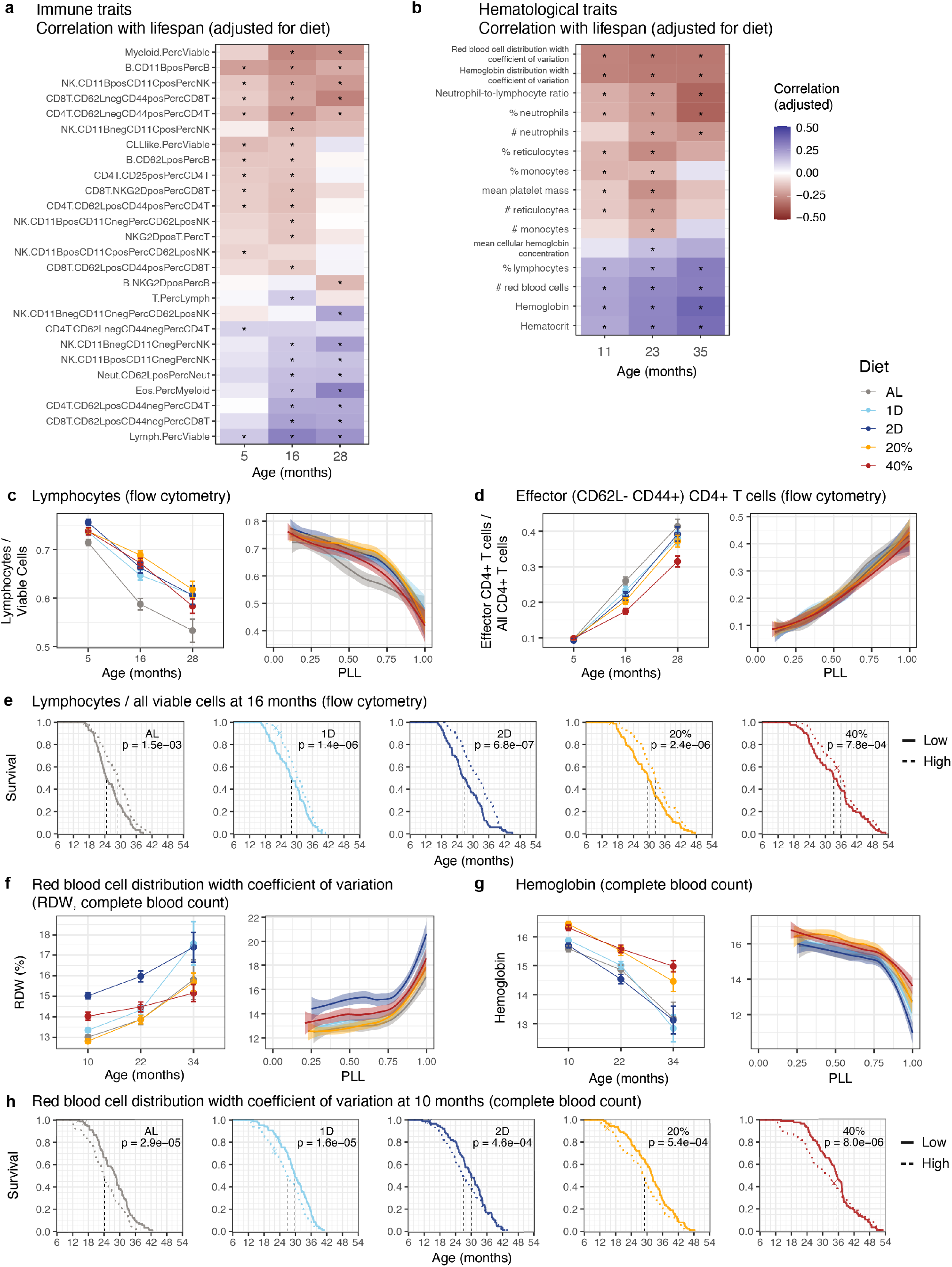
Immune and hematologic traits change with age, respond to diet, and predict lifespan. **a, b,** Heatmaps show the correlation with lifespan for (a) selected traits from the flow cytometry assay and (b) selected traits from the complete blood count assay (y-axis) at ages indicated on the x-axis. Asterisks indicate significant association between the trait and lifespan (FDR < 0.01). Correlations were computed from rank normal scores transformed data and adjusted for diet. **c, d,** Lymphocytes as a proportion of viable cells (c), effector CD4+ T cells as a proportion of all CD4+ T cells (d). **e**, Kaplan-Meier curves for each diet group comparing survival of mice stratified by within-group median lymphocyte proportion at 16 months. Significance was computed by ANOVA of continuous measurements. **f, g,** Red cell distribution width coefficient of variation (RDW, g), and hemoglobin (h) are plotted against age (mean ± 2SE) and against PLL (loess smooth with 95% confidence band). **h**, Kaplan-Meier curves for each diet group comparing survival of mice stratified by within-group median red cell distribution width at 10 months of age. Significance was computed by ANOVA of continuous measurements. Nomenclature for immune cell populations is detailed in **Supplementary Table 7**.

The percent of total circulating lymphocytes in mice at all timepoints was positively associated with lifespan (**Fig. 4a, e**). Cells exhibiting a physiological resting state, such as CD62L^+^CD44^-^ CD4 and CD8 T cells (naïve T cells) and immature NK cells were positively correlated with lifespan, while immune cells displaying evidence of activation or mature phenotypes such as CD62L^-^CD44^+^ CD4 and CD8 T cells (effector T cells) were generally associated with a shortened lifespan (**Fig. 4a**). Designation of immune cell types is detailed in **Supplementary Table 7**.

The major erythroid cell traits (hemoglobin, HGB; hematocrit, HCT; red blood cell count; RBC) decreased with age, while the red blood cell distribution width (RDW, coefficient of variation in volume of red blood cells) and the hemoglobin distribution width (HDW, coefficient of variation in erythrocyte hemoglobin concentration) both increased (**Fig. 4f, g, Extended Data Fig. 9g**). These changes parallel those seen in aging human populations where anemia is a common and significant problem^46,47^. Many of the CBC traits, including RDW and HCT, changed with biological age (PLL scale) with an inflection point and higher rates of change in the last 25% of life. Many of the non-erythroid CBC traits, including neutrophil-to-lymphocyte ratio (NLR) and mean platelet mass (MPM), changed with age but were unaffected by diet (**Extended Data Fig. 9h**). Hemoglobin levels were improved (increased) under CR but not under IF, suggesting a beneficial effect of CR at reducing risk of anemia. Diet response of RDW was distinctive. By 10 months of age, RDW was sharply increased in 2D mice and to a lesser extent in those under 40% CR. The sharp increase in RDW late in life served as an indicator of imminent mortality and partially explained the strong associations between RDW and lifespan. Among the erythroid traits, RDW showed the strongest association with lifespan, and most of the erythroid traits exhibited significant positive associations (HGB, HCT, RBC) or negative associations (RDW, HDW) with lifespan (**Fig. 4b, h**).

### Genetic analysis of lifespan

We obtained whole-genome genotype data for 929 (out of 937) mice and looked at the combined effects of diet and heritability on lifespan. For mice that lived to at least 6 months of age (onset of DR), genetic background explained 23.6% of variation in lifespan (h^2^ = 0.236, 95% bootstrap CI [0.106, 0.360]), while diet explained only 7.4% of variation (**Extended Data Fig. 10a**). As mice aged, heritability declined to 17.1% for mice surviving past 12 months and to 15.9% for mice surviving past 18 months. In parallel, the contribution of diet to lifespan increased with age to 8.4% at 12 months and to 11.4% at 18 months (**Extended Data Fig. 10b, c**). Similar trends of decreasing genetic effect and increasing effects of DR were previously reported for bodyweight^48^.

We carried out genetic mapping analysis of lifespan in the DO mice. We identified a significant quantitative trait locus (QTL; adjusted p < 0.05) for lifespan on chromosome 18 at 21.55Mb (95% support interval [20.44, 24.92]) (**Fig. 5a, Extended Data Fig. 10d**). The DO mice are descended from eight inbred founder strains, and we attributed the QTL effects to the haplotype inherited from the strain CAST/EiJ (CAST) (**Extended Data Fig. 10e**). Among the 164 (out of 929) mice with at least one copy of the CAST haplotype at this locus, lifespan was reduced by an average of 3.7 months (12.5%, p = 6.66e-07). The chromosome 18 QTL effect explained 4.34% of the diet-adjusted variance in lifespan, corresponding to 23.4% of the genetic effect. Despite this, we did not detect any additional significant QTL, and more than 75% of the genetic contribution to lifespan remained unexplained. Kaplan-Meier analysis of lifespan, stratified by the QTL genotype, confirmed that the overall effect of diet was significant in both genotype groups (**Fig. 5b**). However, when the CAST allele was present, only the 40% CR group was significantly different from AL in pairwise comparisons (**Fig. 5b, Supplementary Table 8**). Comparing the QTL effect on lifespan within diets showed that the presence of a CAST allele significantly reduced lifespan for all diet groups except 40% CR (**Fig. 5c**).

**Fig. 5.**
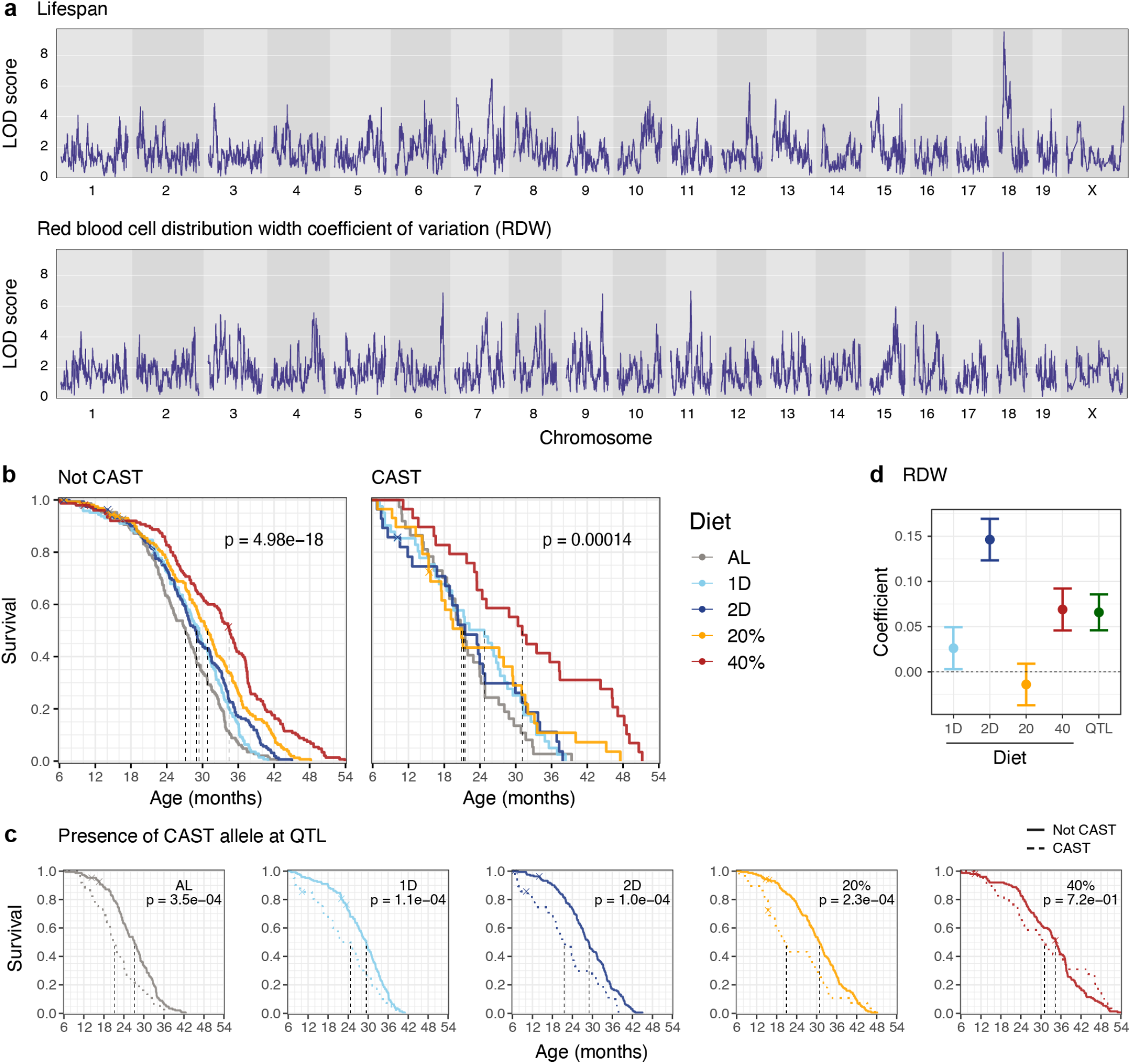
Genetic effects on lifespan in DO mice. **a,** Linkage mapping identifies genome-wide significant association with lifespan and with red blood cell distribution width coefficient of variation (RDW) on mouse chromosome 18. X-axis is genome position and y-axis is log10 likelihood ratio (LOD score). **b**, Kaplan-Meier survival curves by diet group (color) are shown for mice stratified by the presence of a CAST allele at the QTL. Dashed lines indicate median survival time per group. **c**, Kaplan-Meier curves within each diet group compare survival of mice stratified by the presence of a CAST allele at the QTL. Significance is shown as the uncorrected p-value from within-diet linear regression of survival time on the binary genotype. **d**, Estimated regression coefficients show the effects of diet and QTL genotype on RDW. Diets effects are shown as change relative to AL, and genetic effect (green) is response of mice with at least one CAST allele relative to mice with no CAST allele at the chromosome 18 QTL.

Although genetic mapping of physiological traits was not our focus, we identified a QTL for RDW (10 months) that co-localized with the lifespan QTL on mouse chromosome 18 at 21.55Mb (95% support interval [21.06, 21.61]) (**Fig. 5a, Extended Data Fig. 10f**). Presence of the CAST haplotype at this locus was associated with higher RDW (p = 7.08e-11; **Extended Data Fig. 10g**). The effect of the QTL on RDW was similar in magnitude to the effect of 40% CR on RDW (**Fig. 5d**). Variant association mapping of both lifespan and RDW localized the genetic effect (**Extended Data Fig. 10h, i**) to a region that encompasses ∼20 positional candidate genes, including *Garem1, Klhl14, Asxl3, Mep1b, Rnf138, Rnf125,* and a cluster of desmocollin and desmoglein family members (*Dsc* and *Dsg*). The latter have a plausible functional role in maintaining structure and size of red blood cells, but further work is needed to identify a causal gene candidate.

We evaluated RDW as a mediator of the chromosome 18 QTL effect on lifespan. The unconditional effect of the QTL on lifespan was highly significant (QTL → Lifespan, p = 1.85e-7). The QTL effect on lifespan conditioned on RDW was greatly reduced (QTL → Lifespan | RDW, p = 0.00225), and RDW explained 66.0% of the QTL effect on lifespan. These findings suggest that a biological pathway that contributes to variation in mature red blood cell size mediates genetic effects on lifespan. As noted above (**Fig. 4g**), RDW was substantially increased in mice on the 2D IF intervention and to a lesser extent in 40% CR mice. 2D and 40% interventions contributed to reduced lifespan acting through the RDW pathway, and this detrimental effect was offset by the net beneficial effect mediated through other paths (**Fig. 1f**).

## Discussion

In this study, we report the effects of graded levels of CR and IF initiated during adulthood (at 6 months of age) on the health and lifespan of a large cohort of DO mice. We utilized a unique cohort comprising close to 1,000 genetically diverse and genotyped female mice in which we longitudinally profiled 194 traits across 11 categories from the age of 3 months to death. The findings we describe in this cohort have several important implications.

Our results suggest a divergence of the health and longevity effects of DR. Several well-described impacts of DR on metabolic health, such as improved fasting blood glucose, energy expenditure, and circadian oscillations in respiratory quotient, did not predict lifespan within diet groups, as would be expected if these health effects are mediators of the lifespan extension effects of DR. This means that while DR-induced changes in metabolic traits can be beneficial for health, they may not necessarily translate into a significant extension of lifespan. The pleiotropic effects of DR on health span and lifespan may thus be partially non-overlapping, and certain lifespan-extending properties of DR may in fact be detrimental to other aspects of physiological health. This insight has important implications for the choice of biomarkers in human dietary intervention studies, which frequently focus on metabolic health.

In addition, the top predictors of longevity in the DRiDO cohort broadly belong to two categories: body weight/composition and immune/hematological parameters. The top predictors of longevity highlight the importance of body weight resilience during intervention, a high proportion of lymphoid cells (particularly those characterized by naïve/antigen-inexperienced immune cell subsets), a low red cell distribution width, and maintained adiposity late in life. This implies that while some consequences of DR, such as the reduction of myeloid skew with age, directly contribute to its effect on longevity, the ability to withstand certain other impacts of DR, such as those on weight and adiposity, as well as on erythrocyte properties, is beneficial for long-term survival. Importantly, several of these parameters can only be derived from longitudinal monitoring, as performed in this study, highlighting the importance of continuous phenotyping in identifying regulators of DR effects on health and longevity.

Collectively, our study highlights physiological resilience parameters, and in particular the maintenance of body weight, body composition, and immune cell composition over the lifespan, as major biomarkers for longevity and suggests that the pro-longevity effects of DR may be largely uncoupled from its benefit on metabolism.

## Supplementary discussion

### Using large cohorts of DO mice for longevity research

Most studies to date have examined metabolic responses to various forms of DR in male inbred mice, typically C57BL/6J, which have a propensity to develop diet-induced glucose and lipid dysregulation^14,49^. In our study, we focus on female DO mice. These genetically diverse animals present broad phenotypic diversity that can be linked to individual response to DR and increase the translational potential of our findings.

A second notable aspect of our study is the broad range of longitudinal phenotypes measured (194 traits across 11 categories) across the lifetime of the mice. This is important for two reasons: Firstly, we observe that across several traits, including body composition and RDW, association with lifespan varies with the proportion of life lived (PLL) at time of measurement. Secondly, the effects of DR on health and lifespan are complex and pleiotropic, necessitating comprehensive phenotyping of multiple aspects of physiological health. In addition, some of the most important lifespan-associated traits, most notably the ability to maintain body weight, can only be derived from longitudinal monitoring. Despite our best attempts to capture a broad range of physiological traits, our partial correlation network analysis estimates that we are capturing 43% of the effects of DR on lifespan. As discussed in greater detail below, our analysis also indicates that DR affects multiple aspects of physiological health independently, and that such effects, even within groups of related physiological measures, can be both beneficial and detrimental.

### Effects of body weight and composition on longevity

This is the first study that directly compares CR to IF (1 or 2 days fasting), and to report lifespan extension in response to 1D or 2D IF. Mice on CR diets displayed a reduced rate of aging that was proportional to the level of restriction and a corresponding extension of both median and maximal lifespan. IF mice showed extended median lifespan to a lesser degree than CR and did not significantly differ from AL mice in maximum lifespan or rate of aging^50^. It worth noting that the less restrictive IF regimens tested in this study conferred lifespan benefits without significant reduction of net caloric intake.

We confirmed an inverse relationship between early body weight and longevity that progressively decreased in magnitude and reversed later in life^31^. We propose that this reflects a shift in the relative importance of beneficial effects of smaller body size versus higher adiposity as animals age. These observations agree with findings in humans, in which higher weight during childhood is associated with higher morbidity and mortality, and weight loss at older ages is associated with increased frailty and mortality risk^51,52^. Although the relationships between growth, body size, and longevity have been extensively studied, our study unveils several novel interactions between body weight and response to diets, including the lack of response to IF among the heavier mice.

We observed seemingly paradoxical effects of response to DR. Body weight and composition are associated with lifespan in the opposite direction from the intervention effects. DR drives weight loss and reduced adiposity, but mice that maintain more weight and higher adiposity on DR have longer lifespans. The association between lifespan and ability to maintain bodyweight was consistent across all diet groups. The single strongest predictor of lifespan in this study was bodyweight retention in response to stress challenge. We surmise that the ability to maintain bodyweight during periods of stress is an indicator of an individual animal’s resilience. More broadly, DR itself is a stressor and the more resilient animals within an intervention group are better able to maintain bodyweight.

### Metabolic parameters, health, and longevity

Improved metabolic parameters, including weight loss and reduced fasting glucose, represent some of the primary outcomes in human trials of CR, IF and other aging interventions^18–20,22,53–56^. While we see similar changes in metabolic function in the mice, these are not associated with lifespan extension. It seems likely that metabolic changes represent a homeostatic response to reduced caloric intake but that they are not mediating the lifespan extension effects. This raises a question about the interpretation of metabolic response in human trials^23^. The health benefits of reduced adiposity and glucose metabolism for humans are well established, but these may not be good indicators of changes in the rate of aging when analyzed in the context of DR. Intuitively, we may expect that weight loss in response to DR is a sign of the efficacy of the treatment. Rather, our study suggests that individuals who maintain weight on such a regimen may have better long-term outcomes. Specifically, our findings suggest that while DR-induced changes in metabolic traits can be beneficial for health, they may not necessarily translate into a significant extension of lifespan. This poses a challenge when determining the most appropriate primary endpoints for evaluating the effects of interventions in human aging trials. Researchers may need to consider other factors or endpoints when designing and evaluating interventions aimed at promoting healthy aging and longevity. This could include measures related to incidence of chronic disease and mortality and potentially other biological markers associated with functional abilities and resistance to stress.

### Body temperature and lifespan

It has been suggested that lower body temperature (Tb) may play a role in the lifespan-extending effect of CR in rodents^57,58^, non-human primates^59^, and humans^60^. Lowering Tb via genetic manipulation^61^ or changes in ambient temperature^62^ have been reported to promote lifespan independent of energy intake. On the contrary, a study examining the impact of CR on female mice from 28 strains, reported that strains experiencing the most significant drop in Tb were more susceptible to shortened lifespan as compared to those maintaining higher Tb under CR^63^. In this study, we show that Tb generally declines with age and in response to diets. Yet, mice that maintain higher Tb within diet groups had longer lifespan, which seems at odds with the notion that reduced Tb is a mediator of lifespan extension under CR.

### Immune and hematologic traits and lifespan

In contrast to the health and metabolic traits, immune and hematologic traits proved to be powerful predictors of lifespan. The CBC is a routinely used medical diagnostic tool. Decreased hemoglobin levels in aging humans is a strong predictor of all-cause mortality as is an elevated RDW, even in the absence of anemia or other age-related diseases^64,65^. In our DO populations, these traits showed strong positive (HGB, HCT, RBC) and negative (RDW, HDW) associations with lifespan. Ironically, 2D IF and 40% CR extend lifespan while driving important lifespan predictors like RDW in the wrong direction, again pointing to contradictory effects of DR. In other words, DR extends lifespan overall despite having a negative effect on lifespan specifically via pathways associated with RDW.

The response of immune cell populations to DR is complex yet clearly important as an indicator of efficacy of DR. Our findings reveal that DR exacerbates the age-related decline in mature CD11b^+^ NK cells and reduces the level of circulating inflammatory monocytes in a calorie titratable manner, while the increase in percent of circulating B cells with age was completely inhibited only by 40% CR. A recent study indicated that DR modifies trafficking of immune cells between the circulation and tissue sites^66^. Thus, the observed changes in immune cell subsets impacted by diet may have been reflective of differences in tissue residence induced by acute effects of fasting and timing of blood sampling. Nonetheless, after adjusting for diet, changes in circulating immune cell subsets retained their predictive value for longevity.

NK cells are contributors to the innate immune response in that they respond immediately to viral and pathogen challenge in a non-antigen specific context. In some disease settings, mice undergoing caloric restriction have poor outcomes and this may be due to reduced mature NK cell subsets^67–70^. The result in the DO mice suggests that while the NK cell subsets may be exhibiting a less mature phenotype with DR, within the NK cell subsets defined by CD11c and CD11b expression there is an overall shift in frequency to CD62L positive cells with all dietary interventions and this shift is associated with longer lifespan. Since CD62L expression may enhance trafficking of NK cells to sites of viral infection^71^, a relative increase in CD62L by the less mature NK cell subsets may offset potential negative effects of CR on innate NK cell mediated pathogen-directed responses. In general, maintenance of immune cells in a naïve or less activated state correlated well with longevity after adjusting for the effects of dietary interventions.

### Tumors

While tumor incidence was not formally included as part of the health span measures, we did see that DR had an impact on easily observable palpable masses, and distended abdomen. Other studies have shown that DR can impact both tumor incidence and lifespan extension^50,72,73^. It is therefore plausible that the correlation between longevity and immune cell subsets could be at least in part attributed to an effect on tumor development.

A 10% reduction in calories effected by 2D fasting had a greater protective effect on tumor development than the 20% CR intervention. This suggests that caloric intake alone is not the direct cause of tumor control. While the effect of fasting on tumor type development has been demonstrated, our data were not designed to identify the specific neoplastic lesions commonly observed in mice. Therefore, we may have not identified animals carrying less apparent yet lethal cancers.

### Conclusions

One objective of this study was to determine whether IF could be a substitute for CR in animal studies. IF is easier to implement and is a potential alternative to CR for people^21^. IF, as implemented here by withholding food for 1 or 2 days per week does not seem to be as effective at lifespan extension as daily CR, and the heaviest animals appear to derive no benefit from IF. There is no simple direct translation of the 1D and 2D IF interventions between mice and humans. The IF diets in mice result in weekly fluctuation of up to 10% in bodyweight, which resembles “yo-yo dieting” in humans^74,75^. IF and CR diets presented contrasting effects on important health traits. For example, the two CR groups maintained the highest hemoglobin content (HGB, HCT and RBC count), an important health benefit that is not replicated by IF. On the other hand, IF mice maintain more lean mass throughout life. Overall, our findings suggest that IF and CR confer longevity benefits to different degrees and with distinct effects on organismal health.

It is important to consider the balance of beneficial and detrimental effects of different forms of DR. Mice on 40% CR achieved a maximum bodyweight that was only 59.6% of that achieved by AL mice. While these mice are healthy by most measures, we see multiple adverse effects including lower body temperature, food seeking behavior (an indication of hunger), and changes in immune repertoire that could potentially confer susceptibility to infection. These effects in mice may raise concern regarding the benefits of severe restriction for human healthspan. In contrast, 20% CR produced significant lifespan extension, improved health in late life and the impact on both acute and lifetime bodyweight changes is moderate with no loss of adiposity.

Despite the powerful effects of DR in this study, genetic background proved to be the more important factor in determining lifespan. While our study revealed only one significant genetic association with lifespan, it is clear from our heritability analysis that lifespan is regulated by numerous loci with more subtle individual effects, and it is likely that gene-by-gene and gene-by-diet interactions play a role. Our analysis does not account for such gene-by-gene interactions or loci that exert effects only in particular contexts.

It remains unclear which aspects of DR are responsible for lifespan extension, but our study suggests that the pro-longevity effects of DR are largely uncoupled from the benefit on metabolism. Recent studies have attempted to deconvolve the effects of caloric content and timing of feeding, and it appears that both are contributing to lifespan extension^7,8^. Among the DR interventions tested here, 20% CR – with its moderate reduction in caloric intake and consequent daily fasting cycles – may be most easily tolerated by humans. This study provides some of the strongest evidence yet that IF and CR would extend lifespan in humans, but this awaits definitive investigation^76^. Although further work is needed, we suspect that moderate reduction of caloric intake and regular daily feeding and fasting cycles are the key contributing factors to lifespan extension and maximizing the health benefits of DR.

---

**Extended Data Fig. 1.**
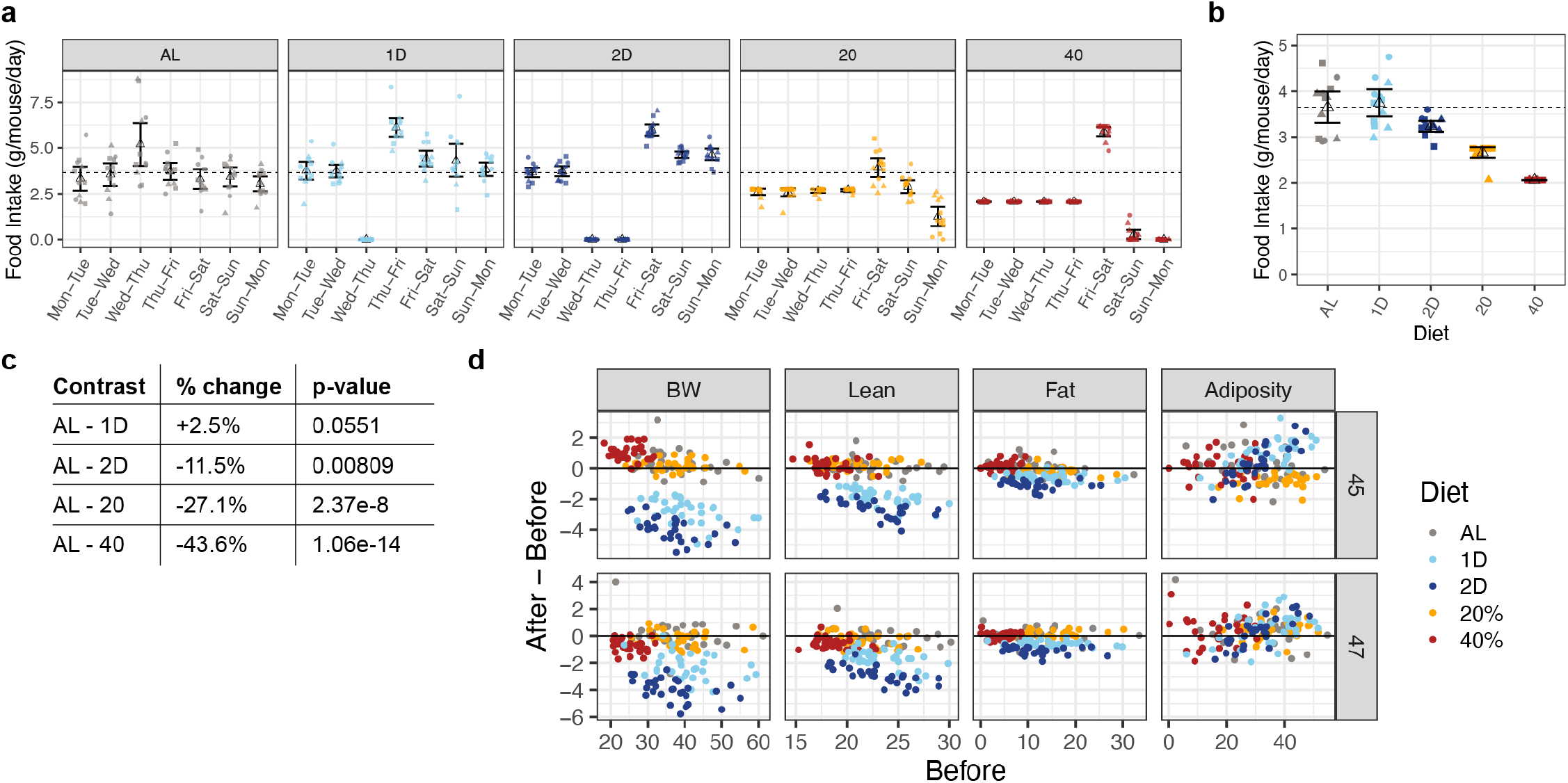
The impact of DR on food consumption. **a,** Food consumption is shown by diet group as daily pen-averages (g/mouse/day) for each of three testing weeks, overlaid by mean ± 2SE. Note that fresh food was provided weekly on Wednesday for AL mice. **b**, Weekly pen-averages (g/mouse/day) food consumption are shown across diets (mean ± 2SE). **c**, Weekly average food consumption in each diet group is summarized as percent difference relative to AL with significance testing by ANOVA. **d**, Changes in body weight and composition before and after fasting at two different testing weeks (45 and 47 weeks). Points represent individual mice at each week of testing. On x-axis: before-fasting bodyweight (g), lean mass (g), fat mass (g), and adiposity (100% x fat mass/total mass). On y-axis: difference between before and after, same units.

**Extended Data Fig. 2.**
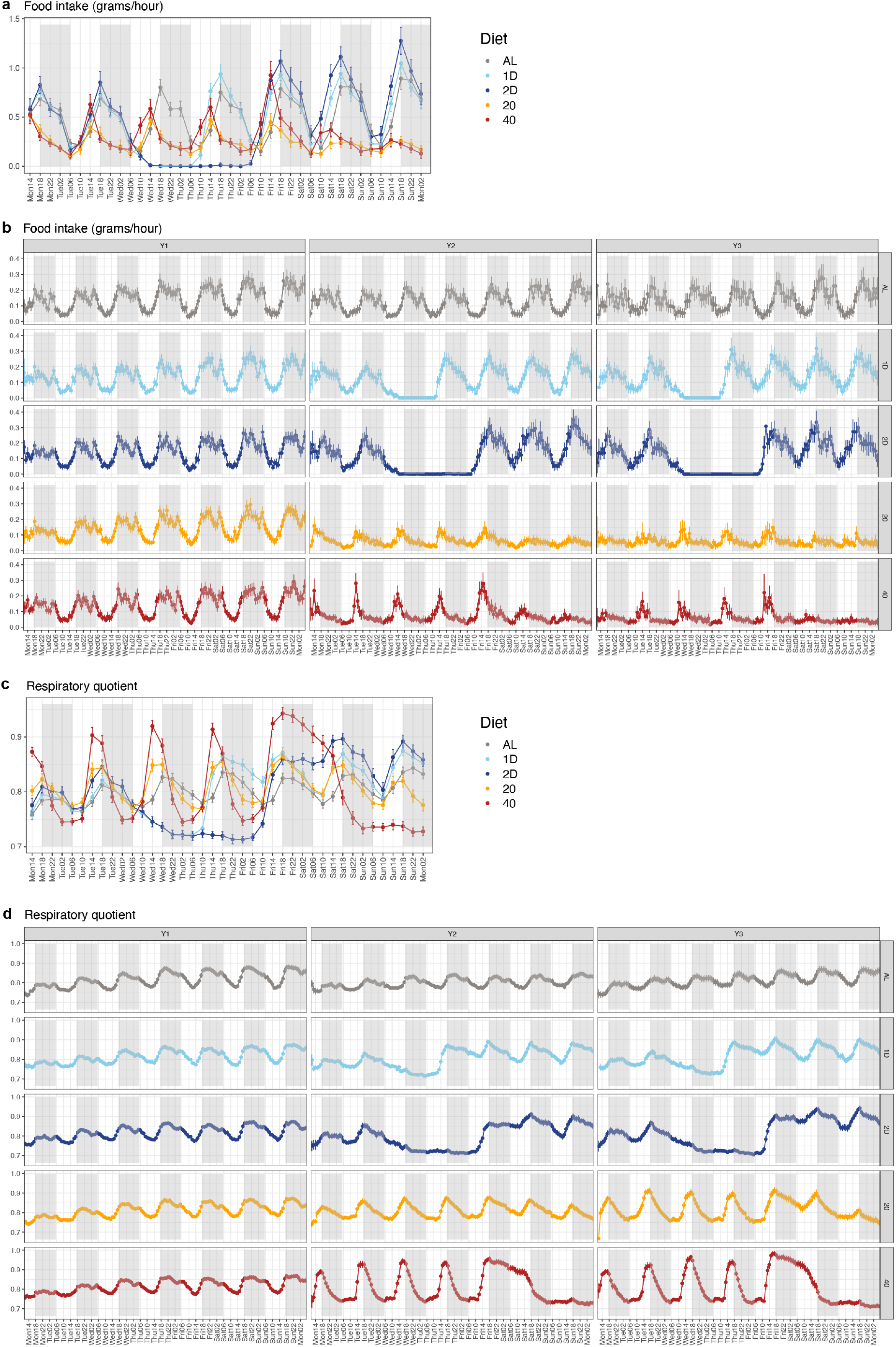
Metabolic consequences of DR. **a,** Food consumption from year 2 metabolic cage data summarized in 4-hour intervals (mean ± 2SE). x-axis labels represent start of 4-hour interval in military time. **b**, Food consumption (grams per hour) in 1-hour intervals (mean ± 2SE). **c,** Respiratory quotient from year 2 metabolic cage data summarized in 4-hour intervals (mean ± 2SE). **d**, Respiratory quotient in 1-hour intervals (mean ± 2SE).

**Extended Data Fig. 3.**
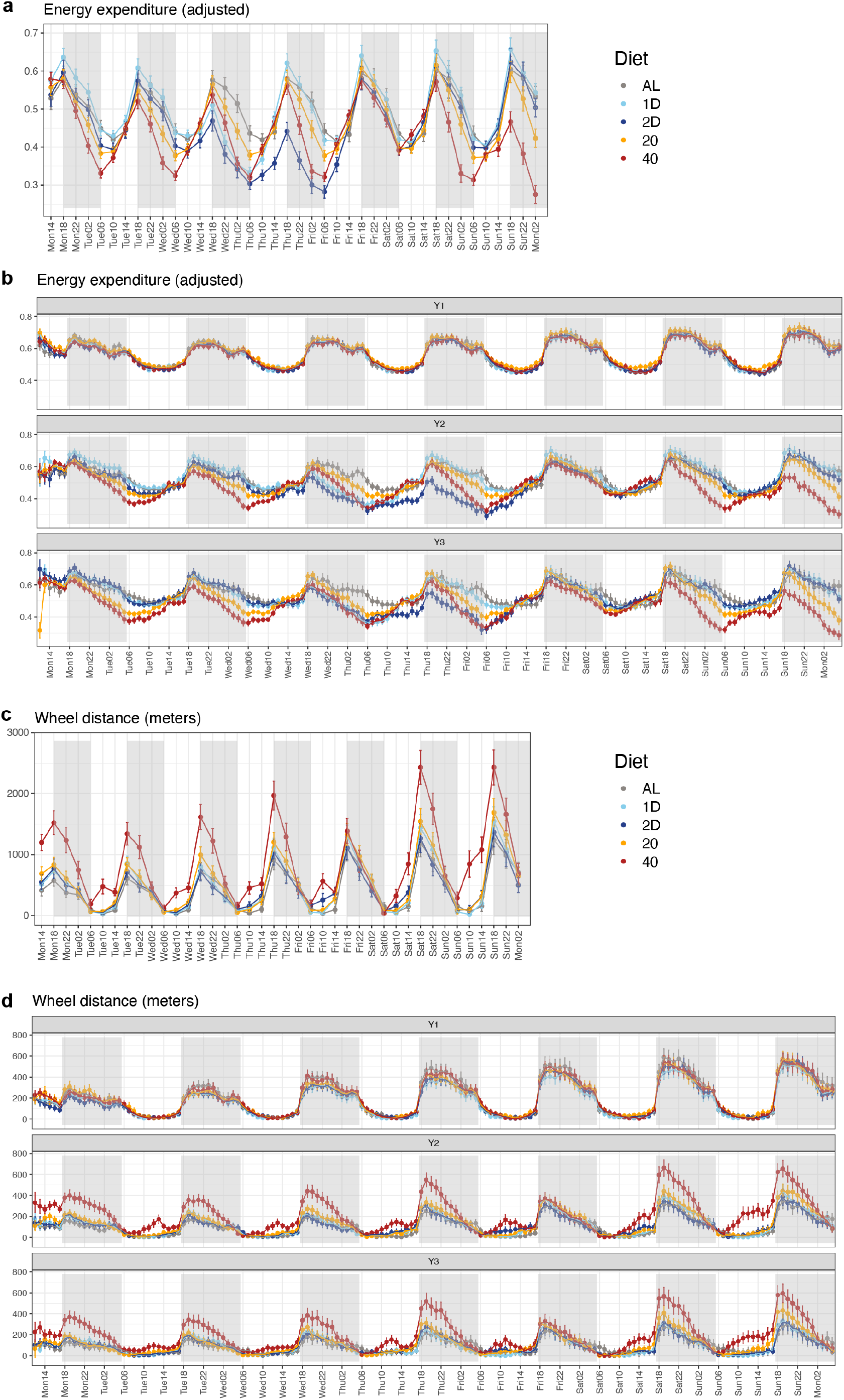
Metabolic consequences of DR. **a**, Energy expenditure (adjusted for bodyweight) from year 2 metabolic cage data summarized in 4-hour intervals (mean ± 2SE). **b**, Energy expenditure in 1-hour intervals (mean ± 2SE). **c,** Distance on running wheel (meters per 4 hours) from year 2 metabolic cage data (mean ± 2SE). **d**, Wheel running (meters per hour) (mean ± 2SE).

**Extended Data Fig. 4.**
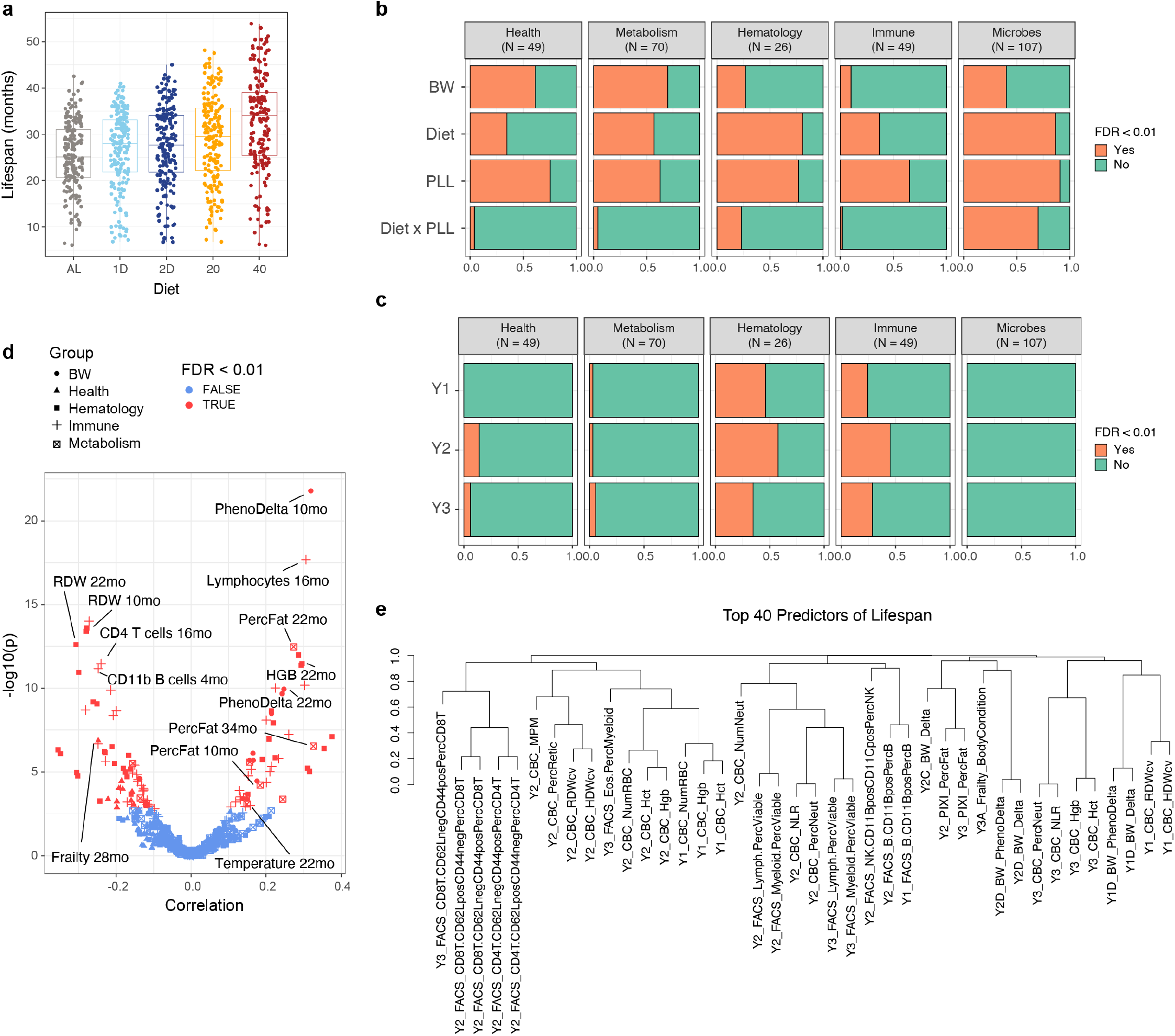
Variables associated with lifespan. **a**, Lifespan variability in mice on different dietary interventions. **b**, Barplots showing the number of traits with significant (FDR < 0.01) associations with bodyweight (BW), diet, biological age (PLL), and diet x PLL interactions. Traits were categorized as health, metabolism, hematology, immune, or microbiome. **c**, Barplots showing the number of significant lifespan associations for traits at each of three ages (designated Y1, Y2, Y3). Ages vary depending on the assay. For traits with multiple measurements in each year, we counted the most significant result. **d**, Volcano plot showing correlations of all assessed traits with lifespan. **e**, Dendrogram shows hierarchical clustering with absolute correlation distance of 40 traits with the most significant lifespan association. Trait names are shown as Year_AssayType_TraitName and detailed in **Supplementary Table 5**.

**Extended Data Fig. 5.**
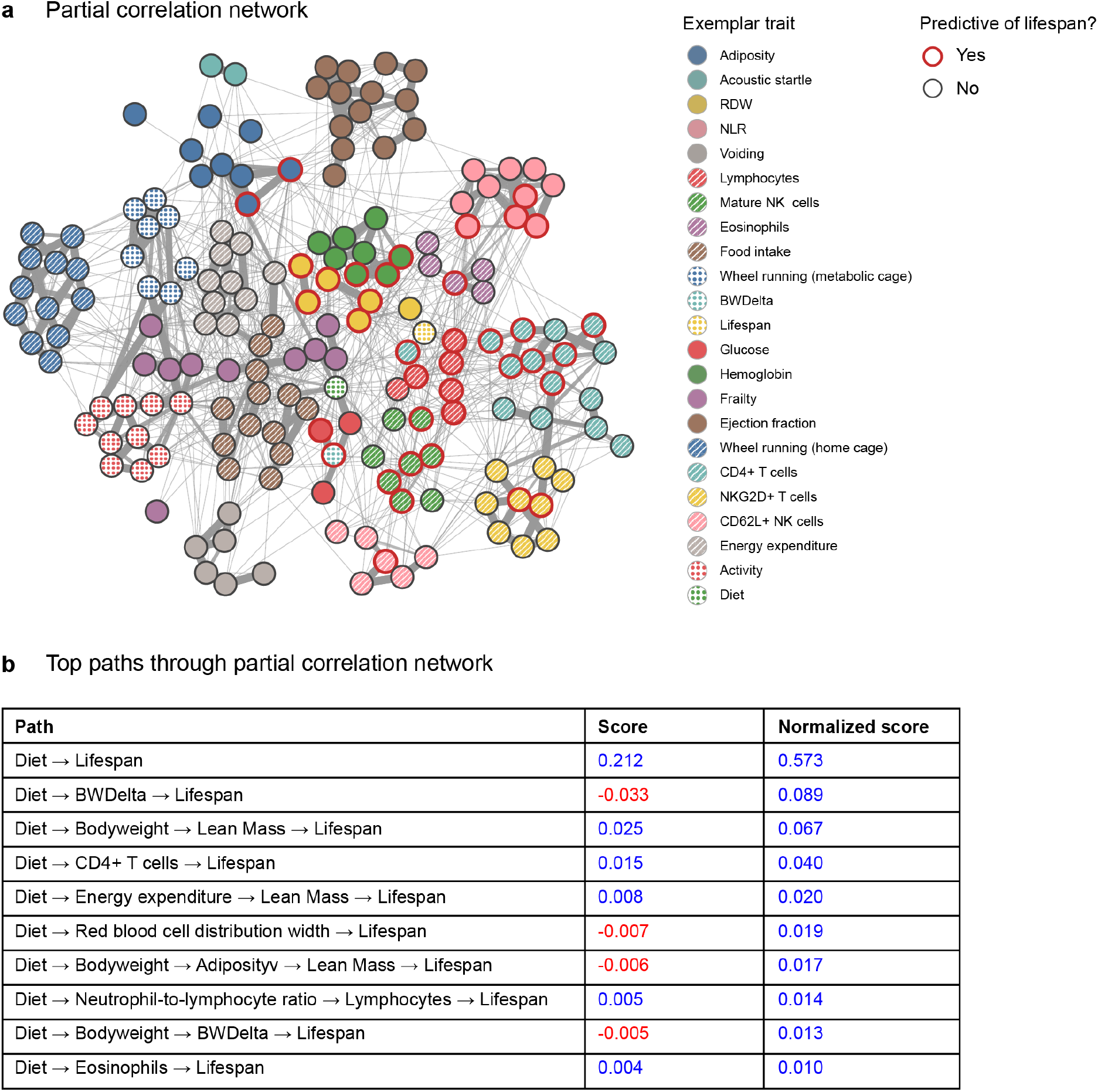
Traits mediating the impact of DR on lifespan. **a**, The partial correlation network of 194 traits. Trait clusters indicated by color with informative naming. Points represent individual traits and red outline indicates traits that are significant predictors of lifespan (FDR < 0.01). Details provided in **Supplementary Table 6**. **b,** The top paths through the partial correlation network linking diet to lifespan. The raw score corresponds to an estimate of the covariance between diet and lifespan mediated via a given path, estimated via covariance decomposition. Paths that show a net negative impact on lifespan are highlighted in red and positive effects paths in blue. The normalized score corresponds to the absolute raw score divided by the sum of all absolute raw scores as a measure of the fraction of diet effects mediated through that path. Note that almost 60% (0.573) of the effect of diet on lifespan is captured in the direct path (Diet -> Lifespan), which represents diet effects on lifespan that are not explained by measured traits.

**Extended Data Fig. 6.**
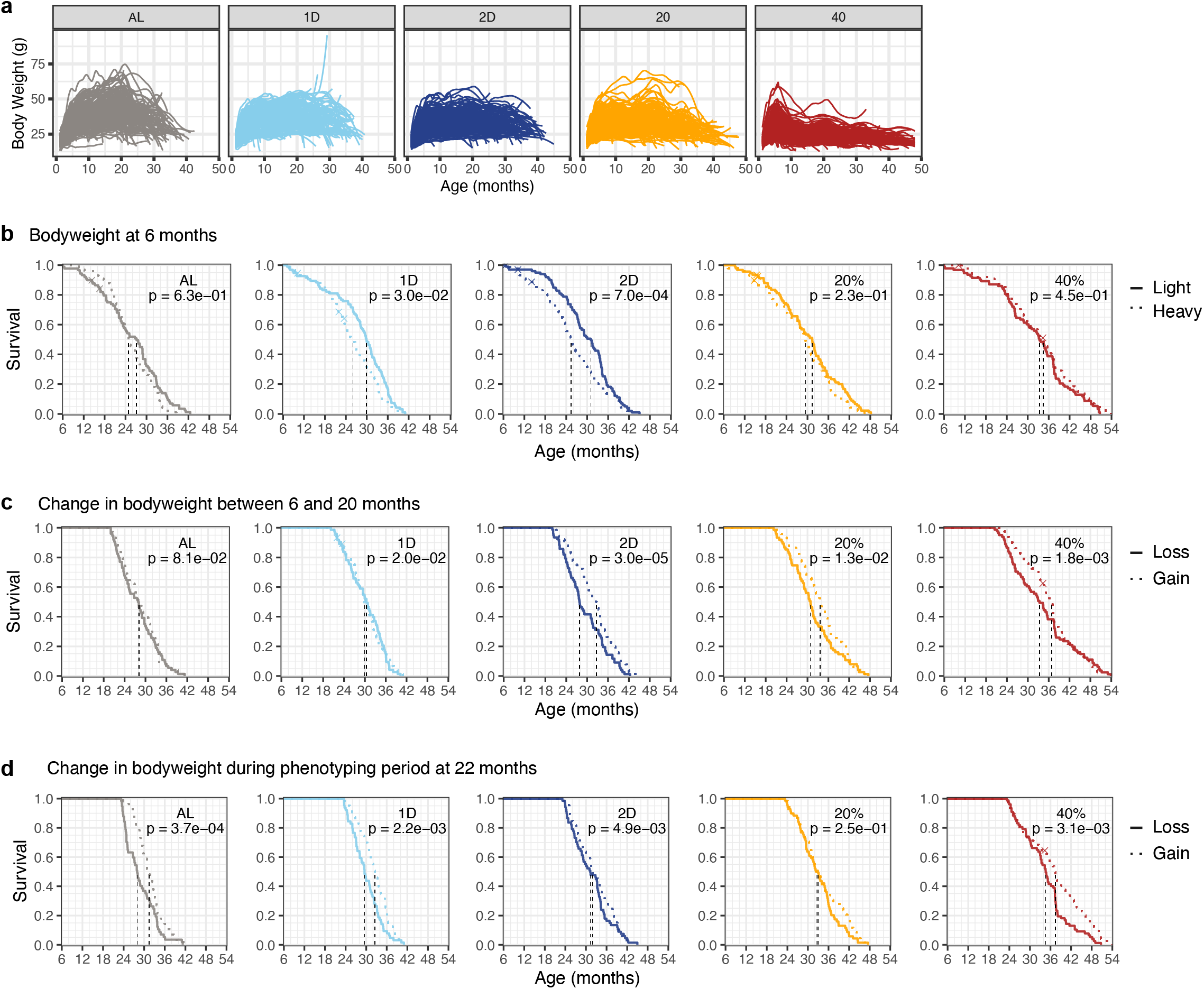
The impact of body weight on lifespan. **a**, Individual bodyweight trajectories by diet group (N = 937 mice). **b-d**, Kaplan-Meier survival curves by diet group (color) are shown for mice below and above median 6-month bodyweight (b), change in body weight between 6 and 20 months (c), and stratified by within-diet median change in bodyweight at 22-23 months of age (d). Significance was assessed on continuous data by ANOVA with adjustment for diet group and p-values are shown as insets.

**Extended Data Fig. 7.**
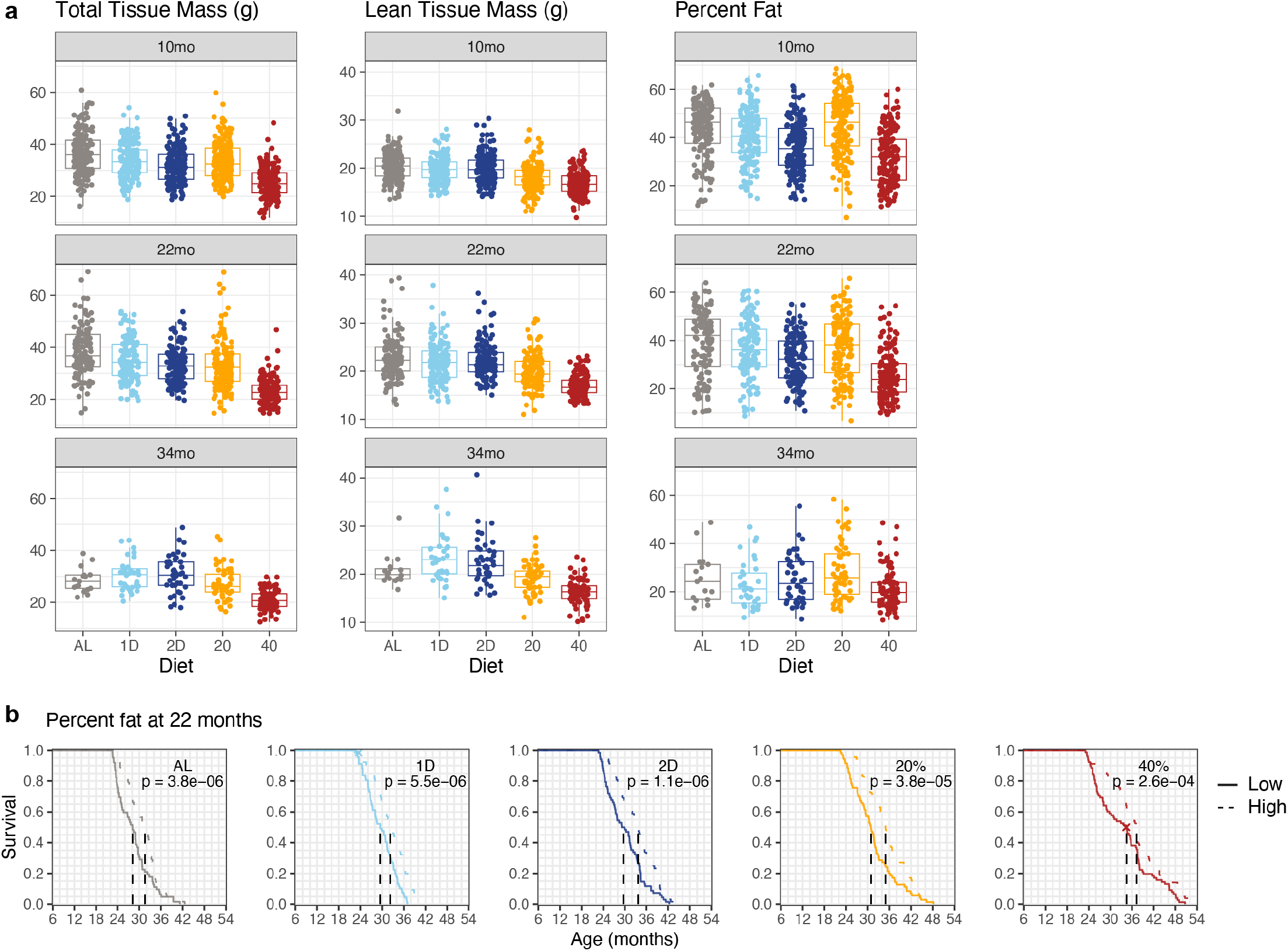
The impact of body composition on lifespan. **a**, Body composition of individual mice at 10, 22, and 34 months of age (N = 895, 689, 241 respectively) showing total tissue mass, lean tissue mass, and percent fat (adiposity = 100% x (1 – LTM/TTM)). All weight measurements in grams (g). **b**, Kaplan-Meier curves within each diet group compare survival of mice stratified by diet-specific median adiposity at 22 months of age. Significance was assessed on continuous data by ANOVA with adjustment for diet group and p-values are shown as insets.

**Extended Data Fig. 8.**
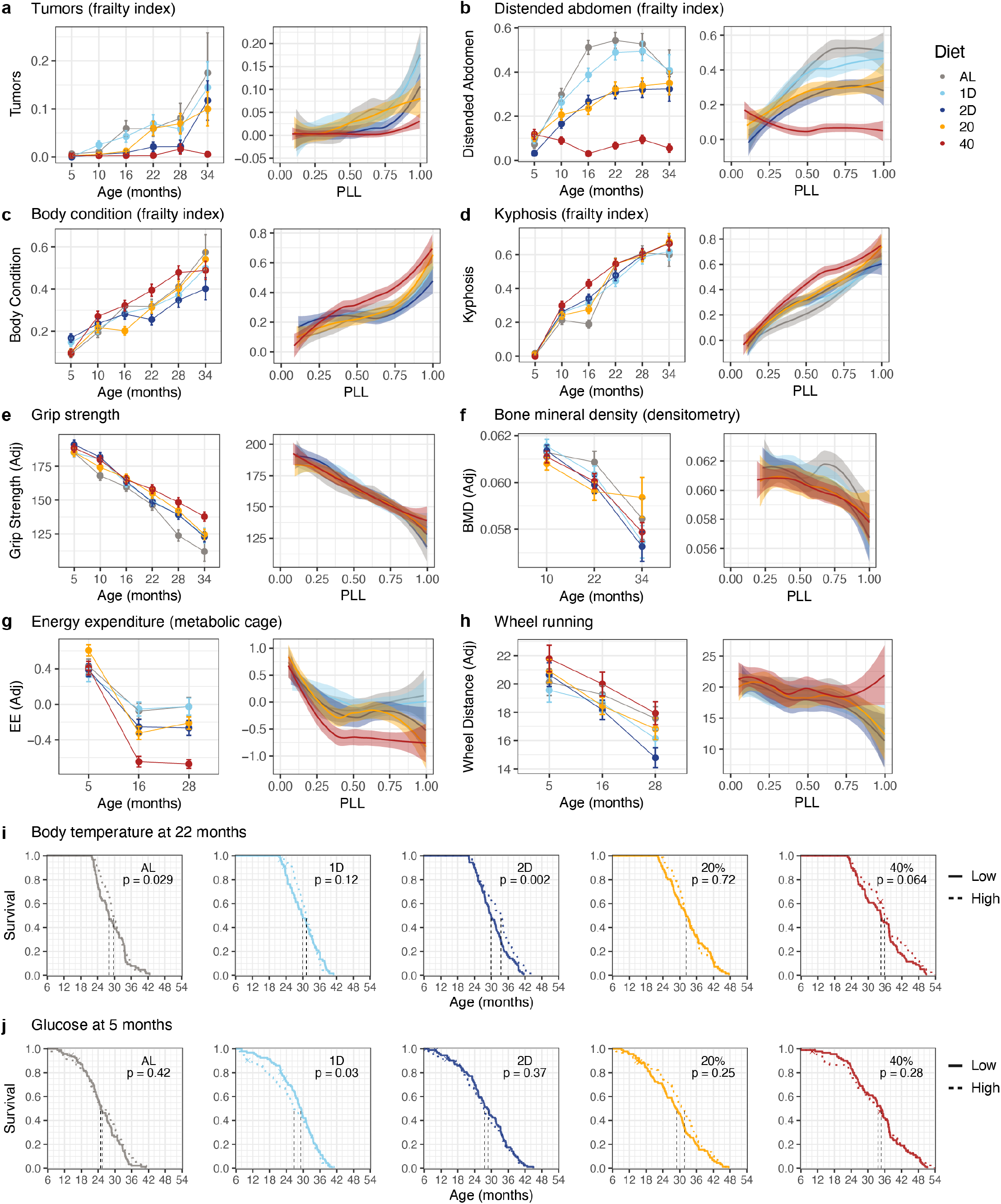
The impact of health parameters on lifespan. **a-h,** Selected traits plotted by chronological age (mean ± 2SE), and by PLL (loess smooth with 95% confidence band). **i, j**, Kaplan-Meier curves for each diet group comparing survival of mice stratified by within-group median trait. Significance was computed by ANOVA of continuous measurements: within-diet median body temperature at 22 months of age (i), and fasting glucose at 5 months of age (j).

**Extended Data Fig. 9.**
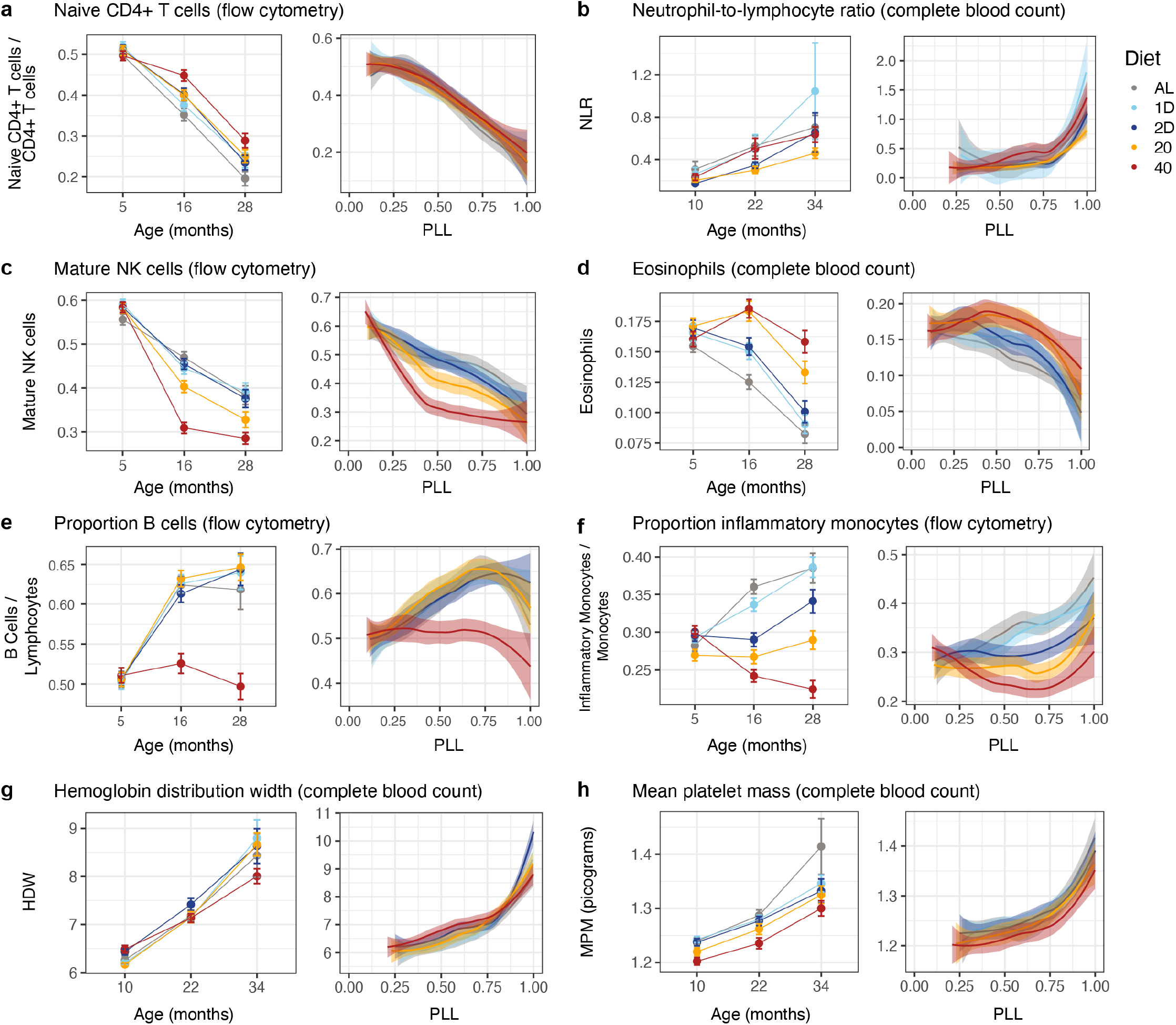
The impact of immune parameters on lifespan. **a-f,** Selected traits from the flow cytometry and complete blood count assays plotted by chronological age (mean ± 1SE), and by PLL (loess smooth with 95% confidence band). **g, h**, Hemoglobin distribution width as coefficient of variation (HDW) (g), and mean platelet mass (MPM, pg) (h) plotted by chronological age (mean ± 2SE), and by PLL (loess smooth with 95% confidence band).

**Extended Data Fig. 10.**
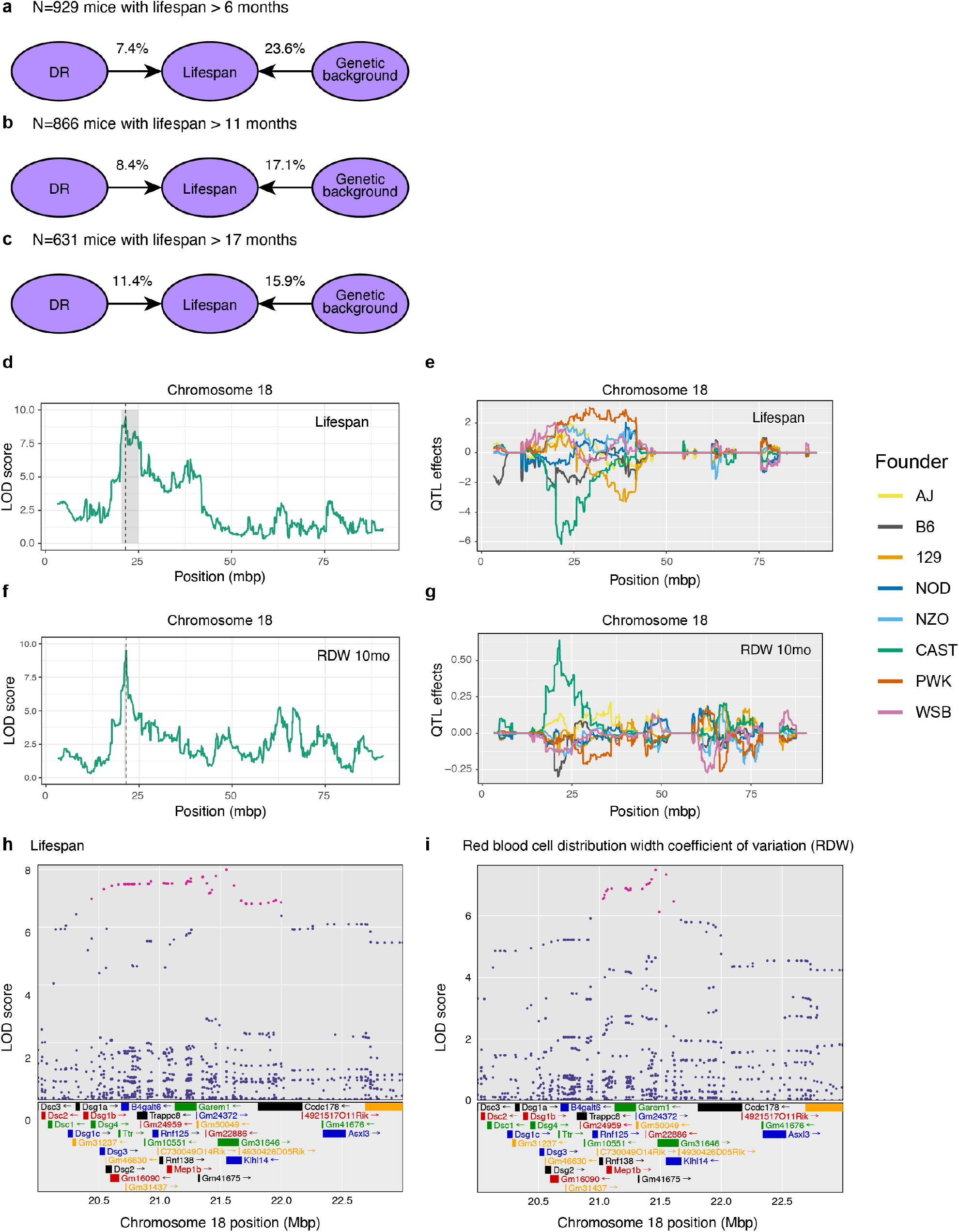
The impact of genetics on lifespan. **a-c,** We estimated the proportion of variance in lifespan explained by diet and genetics for mice that lived to at least (a) 6 months, (b) 11 months, and (c) 17 months of age. **d**, LOD scores from linkage mapping across chromosome 18 of lifespan with diet and 6-month body weight as additive covariates. **e**, Allele effects estimation for lifespan across mouse chromosome 18. **f,** LOD scores from linkage mapping across chromosome 18 of RDW at 10 months with diet and 6-month body weight as additive covariates. **g,** Allele effects estimation for RDW across mouse chromosome 18. **h, i**, Association mapping of single nucleotide variants (SNVs) for lifespan (h) and RDW (i) across the QTL support interval for RDW (21-23Mb). Chromosomal position is shown on the x-axis, SNV LOD score on the y-axis. SNVs with LOD score with 1.5 units of the top scoring SNV are highlighted (purple). Annotated genes and gene models are shown in their approximate positions below.

**Supplementary Table 1:** Pairwise comparison of Diet Effects on Lifespan. Significance (p-value) of all pairwise comparisons (1df) between diet groups from logrank tests. Overall (4df) significance p < 2.2e-16.

|  | 1D | 2D | 20% | 40% |
| --- | --- | --- | --- | --- |
| AL | 0.0433 | 2.65e-4 | 2.92e-8 | 1.39e-19 |
| 1D | -- | 0.0449 | 7.00e-5 | 2.47e-14 |
| 2D | -- | -- | 0.0102 | 4.02e-10 |
| 20% | -- | -- | -- | 1.153e-5 |

**Supplementary Table 2:**
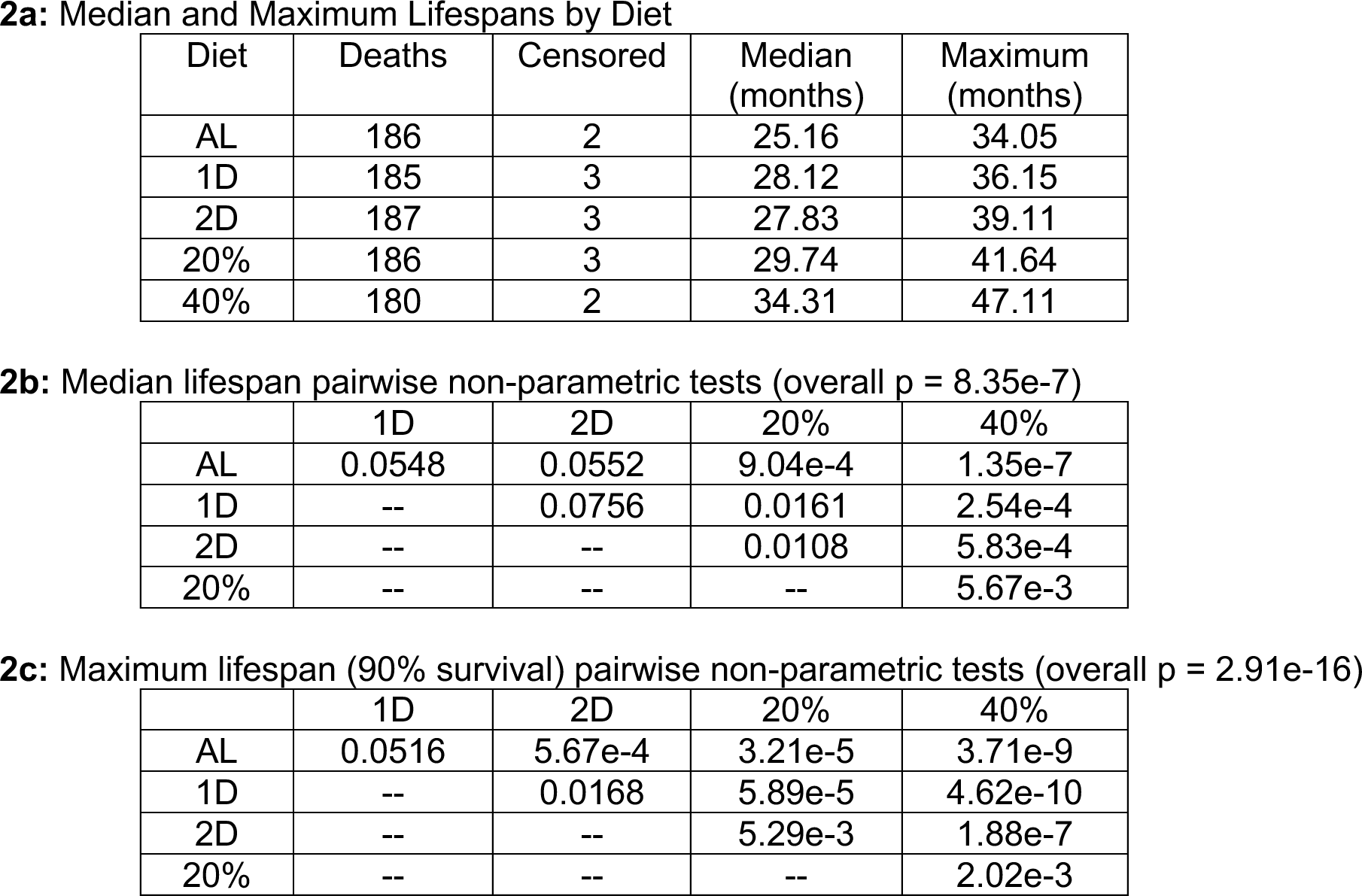
Diet effects on Median and Maximum Lifespan.

2a: Median and Maximum Lifespans by Diet
| Diet | Deaths | Censored | Median (months) | Maximum (months) |
| --- | --- | --- | --- | --- |
| AL | 186 | 2 | 25.16 | 34.05 |
| 1D | 185 | 3 | 28.12 | 36.15 |
| 2D | 187 | 3 | 27.83 | 39.11 |
| 20% | 186 | 3 | 29.74 | 41.64 |
| 40% | 180 | 2 | 34.31 | 47.11 |

2b: Median lifespan pairwise non-parametric tests (overall $p = 8.35\text{e-}7$ )
|  | 1D | 2D | 20% | 40% |
| --- | --- | --- | --- | --- |
| AL | 0.0548 | 0.0552 | 9.04e-4 | 1.35e-7 |
| 1D | -- | 0.0756 | 0.0161 | 2.54e-4 |
| 2D | -- | -- | 0.0108 | 5.83e-4 |
| 20% | -- | -- | -- | 5.67e-3 |

2c: Maximum lifespan (90% survival) pairwise non-parametric tests (overall $p = 2.91\text{e-}16$ )
|  | 1D | 2D | 20% | 40% |
| --- | --- | --- | --- | --- |
| AL | 0.0516 | 5.67e-4 | 3.21e-5 | 3.71e-9 |
| 1D | -- | 0.0168 | 5.89e-5 | 4.62e-10 |
| 2D | -- | -- | 5.29e-3 | 1.88e-7 |
| 20% | -- | -- | -- | 2.02e-3 |

**Supplementary Table 3:** Fitting Gompertz models for DR effects on Lifespan.

3a: Gompertz parameters
| Diet | Baseline Mortality Rate | Mortality Doubling Time | Percent Increase |
| --- | --- | --- | --- |
| AL | 0.00660 | 5.17 | - |
| 1D | 0.00503 | 5.05 | -2.22 |
| 2D | 0.00580 | 5.71 | 10.6 |
| 20% | 0.00625 | 6.53 | 26.4 |
| 40% | 0.00522 | 7.39 | 41.0 |

|  | 1D | 2D | 20% | 40% |
| --- | --- | --- | --- | --- |
| AL | 0.823 | 0.316 | 0.0212 | 0.000797 |
| 1D | -- | 0.229 | 0.0132 | 0.000455 |
| 2D | -- | -- | 0.193 | 0.0184 |
| 20% | -- | -- | -- | 0.292 |

**Supplementary Table 4: Diet, Age and Bodyweight effect on Traits**

Filename: Supplemental_Table4_DietAgeBW.csv

**Supplementary Table 5: Lifespan x trait correlations**

Filename: Supplemental_Table5_SurvCor.csv

**Supplementary Table 6: Network analysis details**

Filename: Supplemental_Table6_NetworkDetails.xlsx

**Supplementary Table 7: Nomenclature for FACS Traits**

Filename: Supplemental_Table7_FACSNames.xlsx

**Supplementary Table 8:** Diet Effects on Lifespan Stratified by Bodyweight at 6 months.

|  | 1D | 2D | 20% | 40% |
| --- | --- | --- | --- | --- |
| AL | 0.185 | 0.00546 | 2.95e-04 | 2.21e-08 |
| 1D | -- | 0.143 | 0.00978 | 1.78e-05 |
| 2D | -- | -- | 0.113 | 0.0101 |
| 20% | -- | -- | -- | 0.0337 |

**8b: Logrank pairwise tests Heavy mice (overall p < 1.20e-15).**
|  | 1D | 2D | 20% | 40% |
| --- | --- | --- | --- | --- |
| AL | 0.238 | 0.0598 | 1.43e-05 | 3.68e-14 |
| 1D | -- | 0.264 | 0.00125 | 4.11e-11 |
| 2D | -- | -- | 0.0201 | 1.32e-08 |
| 20% | -- | -- | -- | 4.50e-05 |

**8c: Median and maximum lifespan by early BW**
| Diet | Median<br>Light/Heavy | p value<br>(7.63e-4) | Max (90%)<br>Light/Heavy | p value<br>(0.167) |
| --- | --- | --- | --- | --- |
| AL | 27.1 / 24.8 | 0.770 | 36.1 / 33.7 | 0.0112 |
| 1D | 29.9 / 26.0 | 0.00141 | 36.8 / 34.6 | 0.125 |
| 2D | 31.1 / 25.4 | 0.00383 | 39.3 / 38.6 | 0.471 |
| 20% | 31.4 / 29.4 | 0.384 | 42.6 / 41.1 | 0.325 |
| 40% | 33.4 / 34.4 | 0.657 | 46.2 / 47.5 | 0.802 |

**Supplementary Table 9:** Diet Effects on Lifespan Stratified by Chromosome 18 QTL.

|  | 1D | 2D | 20% | 40% |
| --- | --- | --- | --- | --- |
| AL | 0.0776 | 0.00161 | 2.08e-07 | 2.22e-15 |
| 1D | -- | 0.0759 | 0.000174 | 2.90e-11 |
| 2D | -- | -- | 0.0113 | 7.46e-08 |
| 20% | -- | -- | -- | 0.000493 |

**9b: All pairwise log-rank tests for mice with CAST allele**
|  | 1D | 2D | 20% | 40% |
| --- | --- | --- | --- | --- |
| AL | 0.326 | 0.439 | 0.171 | 2.97e-05 |
| 1D | -- | 0.983 | 0.472 | 0.00030 |
| 2D | -- | -- | 0.610 | 0.00105 |
| 20% | -- | -- | -- | 0.00477 |

*Companion manuscript*: Litichevskiy, L., Considine, M., Gill, J., Shandar, V., Cox, T. O., Descamps, H.C., Wright, K. M., Amses, K. R., Dohnalová, L., Liou, M. J., Tetlak, M., Galindo-Fiallos, M. R., Wong, A. C., Lundgren, P., Kim, J., Uhr, G. T., Rahman, R. J., Mason, S., Merenstein, C., Bushman, F. D., Raj, A., Harding, F., Chen, Z., Prateek, G. V., Mullis, M., Deighan, A. G., Robinson, L., Tanes, C., Bittinger, K., Chakraborty, M., Bhatt, A. S., Li, H., Barnett, I., Davenport, E. R., Broman, K. W., Cohen, R. L., Botstein, D., Freund, A., Di Francesco, A., Churchill, G. A., Li, M., Thaiss, C.A. Interactions between the gut microbiome, dietary restriction, and aging in genetically diverse mice.

## Methods

### Mice

We enrolled 960 female DO mice in 12 waves, corresponding to birth cohorts from generations 22 to 24 and 26 to 28 with 80 first parity and 80 second parity animals from each generation born ∼3 weeks apart. Just one female mouse per litter was enrolled in the study. The first cohort entered the study in March 2016 and the study was fully populated in November 2017. This schedule was designed to make efficient use of our phenotyping capacity and minimize the potential for seasonal confounding. Study size was determined to detect a 10% change in mean lifespan between intervention groups with allowance for some loss of animals due to non-age-related events. The study included only female mice. Mice were assigned to housing groups of 8 animals per pen in large format wean boxes. Cages were positive pressure ventilated with air temperature of 76 to 78F. Environmental enrichments were provided including nestlets, biotubes, and gnawing blocks. Mice were randomized by housing group to one of 5 dietary interventions. Of the 960 mice that were entered into the study, 937 mice were alive when interventions were initiated at 6 months of age and only these mice are included in our analysis. All procedures used in the study were reviewed and approved under Jackson Lab IACUC protocol #06005.

### Dietary Restriction

Dietary restriction (DR) was implemented by controlling the timing and amount of food provided to mice. Feeding schedules for DR were started at 6 months of age. All mice were fed a standard mouse chow diet (5K0G, LabDiet). The *ad libitum* (AL) feeding group was provided with unlimited access to food and water. The intermittent fasting (IF) mice were provided unlimited access to food and water. On Wednesday of each week at 3pm, IF mice are placed in clean cages and food was withheld for the next 24 or 48 hours for the 1D and 2D groups, respectively. Calorie restricted (CR) mice were provided with unlimited access to water and measured amounts of food daily at ∼3pm, 2.75g/mouse/day for 20% CR, and 2.06g/mouse/day for 40% CR. These amounts were based on AL consumption of 3.43g/mouse/day that we estimated based on historical feeding data from DO mice. Mice were co-housed with up to 8 mice per pen. Co-housing is standard practice for CR studies^15^; competition for food was minimized by placing food directly into the bottom of the cage, allowing individual mice to ‘grab’ a pellet and isolate while they eat. CR mice were provided with 3 days of food on Friday afternoon, which resulted in weekly periods of feasting followed by a period of food deprivation of approximately for 1 day for the 20% CR mice and 2 days for the 40% CR mice, comparable to the IF fasting periods.

### Food Intake (160 mouse independent cohort)

To obtain an accurate estimate of food intake and changes in bodyweight in response to weekly fasting cycles, we set up an independent cohort of 160 female DO mice. Mice were placed on the same DR protocols as in the main study. Food was weighed daily for a period of one week when mice were 30, 36 and 43 weeks of age. Food intake data were normalized to units of g/mouse/day and presented as daily and weekly averages across timepoints by diet. Body composition was determined at 43 and 45 weeks of age by non-imaging nuclear magnetic resonance (NMR) using the Echo MRI (Houston, TX) instrument, a nuclear magnetic resonance device (NMR) with a 5-gauss magnet that is adapted to small animal studies. NMR data were used to detect changes in body weight and composition before and after fasting. Values of bodyweight, lean mass, fat mass, and adiposity (100% x fat mass/total mass) pre-fasting were co-plotted with the difference between before and after fasting (Friday to Monday for AL and CR; Tuesday to Thursday for 1D IF; Tuesday to Friday for 2D IF).

### Phenotyping

We carried out three cycles of health assessments of mice at early, middle, and late life. These assessments included a 7-day metabolic cage run at ∼16 weeks, 62, and 114 weeks of age; blood collection for flow cytometry analysis (FACS) at 24, 71, and 122 weeks; rotarod, body composition, echocardiogram, acoustic startle, bladder function, free wheel running, and a blood collection for whole blood analysis (CBC) at ∼44, 96, and 144 weeks of age. In addition, body weights were recorded weekly and manual frailty and grip strength assessments were done at 6-month intervals. All assays were conducted at The Jackson Laboratory following standard operating procedures.

#### Bodyweight

Mice were weighed weekly throughout their lives, resulting in over 100,000 values longitudinally collected for the 937 mice. Body weights were analyzed after local polynomial regression fitting within mouse (i.e., loess smoothing).

#### Frailty, grip strength, and body temperature

We applied a modified version of the clinically relevant frailty index^37^, which was calculated as the average of 31 traits that are indicators of age-associated deficits and health deterioration. Each trait was scored 0, 0.5, or 1 scale where 0 indicated the absence of the deficit; 0.5 indicated mild deficit; and 1 indicated severe deficit. Measurements were taken at baseline (5-months) and repeated approximately every 6 months. Simple averaging yielded a raw frailty index score between 0 and 1 for each mouse. Frailty scores were adjusted by estimating batch and experimenter effects as random factors that were subtracted from raw frailty score values prior to statistical analysis.

#### Body composition

The LUNAR PIXImus II densitometer was used to collect bone density and body composition (including fat and muscle tissue). Mice were anesthetized and individually placed on a disposable plastic tray that was then placed onto the exposure platform of the PIXImus. The process to acquire a single scan lasts approximately 4 minutes; data can be manipulated subsequently to obtain specific regions of interest. Measurements were taken at ∼44, 96, and 144 weeks of age.

#### Immune cell profiling by flow cytometry

Peripheral blood samples were analyzed by flow cytometry to determine the frequency of major circulating immune cell subsets. Analysis was performed prior to the start of dietary interventions at 5 months, then at 16 and 24 months of age. These timepoints corresponded with 11 and 19 months of dietary intervention. Red blood cells in PBL samples were lysed and the samples were washed in FACS buffer (Mitenyi 130-091-222). Cells were resuspended in 25 ul FACS buffer with 0.5% BSA (Miltenyi cat# 130-091-222 with 130-091-376). Antibodies including Fc block (clone 2.42, Leinco Technologies) were added and incubated for 30 minutes at 4°C. Labeled cells were washed and DAPI was added prior to analysis on an LSRII (BD Bioscience). The antibody cocktail contained CD11c FITC, Clone N418 (Cat# 35-0114-U100 Tonbo Biosciences (1:100)); NKG2D (CD314) PE, Clone CX5 (Cat# 558403 BD Biosciences (1:80)); CD3e PE-CF594, clone 145-2C11 (Cat# 562289 BD Biosciences (1:40)); CD19 BB700, clone 1D3 (Cat# 566411 BD Biosciences (1:40)); CD62L PE-Cy7, clone MEL-14, (Cat# 60-0621-U100 Tonbo Biosciences (1:100)); CD25 APC, Clone PC61 (Cat# 102012 Biolegend (1:80)); CD44 APC-Cy7, Clone IM7 (Cat# 25-0441-U100 Tonbo Biosciences (1:40)); Ly6G BV421, Clone 1A8, (Cat# 562737 BD Biosciences (1:80); CD4 BV570, Clone RM4-5 (Cat# 100542 Biolegend (1:40)); CD11b BV650, Clone M1/70 (Cat# 563402 BD Biosciences (1:160)); CD45R/B220 BUV496 (Clone RA3-6B2, Cat# 564662 BD Biosciences (1:20)); Fc Block, Clone 2.4G2 (Cat C247 Leinco Technologies (1:100)).

Due to the outbred nature of these mice, flow cytometric markers were limited, and T cell subsets were generally assigned as naïve and non-naïve by the presence of CD62L and CD44 (see **Supplementary Table 7** for immune cell subtype designations). NKG2D positive cells were enumerated and may represent memory T cells that accumulate after immune responses^77^. Due to limitations in flow cytometric markers that identify NK cells and their subsets in the mouse strains contributing to the outbred DO mouse line, NK cells were defined as non-T non-B lymphocytes expressing NKG2D. Within this population, CD11c and CD11b were used to generally define maturation subsets. CD11b expression marks more mature NK cells and CD11c is reduced on the least mature NK subset^78^.

#### Glucose

At the flow cytometry blood collections at 16, 62, and 114 weeks, mice were fasted for 4 hours and glucose was measured using the OneTouch Ultra2 glucose meter from LifeScan along with OneTouch Ultra test strips. At each of the CBC blood collections at 24, 71, and 122 weeks, non-fasted glucose was measured using the glucose meter.

#### Echocardiogram

Ultrasonography was performed using a VisualSonics Inc. (VSI) Vevo 770/2100 high-frequency ultrasound with 30 and 40 MHz probes. Echocardiography uses pulsed Doppler sonography, applied through the ultrasound probe, to measure blood flow rates and volumes.

#### Metabolic monitoring cages

Mice were individually housed for seven days in a metabolic cage (Sable Systems International) and activity, feeding, and respiration were tracked. Feeding protocols for dietary intervention were maintained. Metabolic cage data were used to assess metabolism, energy expenditure, and activity of mice in Y1, Y2, and Y3. Animal-level data were cleaned to remove outliers and instrument failure and summarized as cumulative (Food, RQ) or median across 5-minute intervals (RQ, EE). Mean and SD were computed at 4-hour and 1-hour intervals. Annual summaries across animals were shown as mean ± 2SE across timepoint intervals.

#### Complete blood count (CBC)

Blood samples were run on the Siemens ADVIA 2120 hematology analyzer with mouse-specific software as described previously^79^.

#### Acoustic Startle

Startle response was measured in rodents using automated startle chambers, in which a mouse was placed in a clear, acrylic tube attached to a highly sensitive platform that is calibrated to track their startle reflex while being exposed to a series of stimuli at varying decibels and times. Mice were initially exposed to white noise from an overhead speaker, which transitions to a series of randomized, computer-generated stimuli ranging in volume from 70 to 120 decibels at 40 msec in duration and an interval of 9-22 seconds. The test runs for approximately 30 minutes.

#### Rotarod

We used the Ugo-Basile rotarod, which has 5 lanes evenly spaced along a motorized horizontal rotating rod, allowing for up to five mice to be tested simultaneously. Below each lane is a platform equipped with a trip plate that records the latency for each mouse to fall. At the beginning of the session, mice were placed on the rod, which begins rotating at 4rpm, slowly increasing to a maximum of 40rpm, over 300 seconds. Mice were given three consecutive trials. We reported the mean latency (time to fall) and the slope of latencies across trials, as well as the number of trials with no falls and number of trails with immediate falls. In case a mouse did not cooperate with the test, trails were recorded as missing.

#### Voiding assay

Cages were prepped by cutting a piece of cosmos blotting paper, 360gsm, to standard duplex cage dimensions. Shavings were removed from a clean cage, and the paper was taped to the bottom of the cage. Food was provided during this test; however, water was removed to prevent possible leaking onto the blotting paper. Mice were individually housed in a prepared cage for 4 hours. At the end of the trial, mice were returned their original housing units, and papers were removed and dried for 2-4 hours, before being individually bagged. Papers were shipped to Beth Israel Deaconess Medical Center where they were scanned with UV light to image and quantify the void spots.

#### Home cage wheel running

Free wheel running data were collected at ∼44, 96, and 144 weeks of age. Mice were individually housed for a minimum of 36 hours in a special cage suited to house the Med Associate low profile running wheel with a wireless transmitter. Food hopper was removed to allow for seamless movement of the wheel, and food was placed on the cage floor. The 15.5-cm-diameter plastic wheel sits at an angle on an electronic base which tracks the revolutions. The battery powered base allows for continuous monitoring of data, which is then wirelessly transmitted, in 30 second intervals, to a local computer.

#### Lifespan

Research staff regularly evaluated mice for pre-specified clinical symptomology: palpable hypothermia, responsiveness to stimuli, ability to eat or drink, dermatitis, tumors, abdominal distention, mobility, eye conditions (e.g., corneal ulcers), malocclusion, trauma, and wounds of aggression. If mice met the criteria for observation in any of these categories, veterinary staff were contacted. If the clinical team determined a mouse to be palpably hypothermic and unresponsive, unable to eat or drink, and/or met protocol criteria for severe dermatitis, tumors, and/or fight wounds, preemptive euthanasia was performed to prevent suffering; otherwise, the veterinary staff provided treatment. Both mice euthanized or found dead were represented as deaths in the survival curves. Mice euthanized due to injuries unrelated to imminent death were treated as censored (we recorded a total of 13 censoring events).

### Data preparation and analysis

Cloud-based research management software (Climb by Rockstep, Inc.) was used to track animals, schedule testing, and provide a stable repository for primary data collection. Data were regularly reviewed by a statistical analyst during the study for anomalies. Initial data quality control included identifying and resolving equipment mis-calibration, mislabeled animals, and technically impossible values. If we could not manually correct these using laboratory records, they were removed. Quantitative assays including body weight and temperature were explored for outliers. Quantitative traits other than bodyweights were corrected for batch effects. To quantify batch effects, we fit a fully random effects linear mixed model conditioning on diet, bodyweight at test, and age. We adjusted trait values by subtracting the batch model coefficients. Lifespan data were recorded in days but are presented in months (30.4 days per month) for ease of interpretation. All analyses were performed using R version 4.2.2 and RStudio version 2022.12.0+353. The full statistical analysis code is publicly available, see “Data Availability”.

#### Survival Analysis

We performed survival analysis to compare lifespan outcomes for the five study groups. We plotted Kaplan-Meier survival curves and tested the equality of survival distributions across diet groups using log rank tests using an overall test (4df) and all pairwise comparisons between diets. p-values are reported with no multiple testing adjustment, and we considered a comparison to be significant if p < 0.01. We estimated median and maximum lifespan by diet group with 95% confidence intervals and percent change relative to AL^28^. Mortality doubling times were estimated from a Gompertz log-linear hazard model with 95% CI and percent change relative to AL (flexsurv R package v2.2.2).

#### Longitudinal trait analysis

For traits collected annually or biannually, we were able to explore hypothesized direct and indirect relationships with diet, body weight, and age. We used generalized additive mixed models (GAMMs) with a gaussian/identity link to analyze these effects by fitting a series of nonlinear relationships between trait response and covariates resulting in 970 fits (194 traits x 5 nested models). GAMMs for a combination of fixed and random effects (formulae below) on trait response pre-adjusted for batch were fit using the gam() function from the mgcv package in R, with the newton optimizer and default control parameters. Age was rescaled to proportion of life lived (PLL = age at test / lifespan). The PLL scale removes artifacts due to survivorship bias across groups with different lifespans. all continuous variable except PLL were rank normal scores transformed prior to model fitting.

1) rankZ(*T*) ∼ D + s(BW) + s(PLL) + (1|ID)
2) rankZ(*T*) ∼ D + s(PLL) + (1|ID)
3) rankZ(*T*) ∼ D + s(BW) + (1|ID)
4) rankZ(*T*) ∼ s(BW) + s(PLL) + (1|ID)
5) rankZ(*T*) ∼ D + s(BW) + s(PLL|Diet) + (1|ID)

where *T* is trait, *D* is dietary assignment, *BW* is body weight at date closest to *T’s* collection date, PLL is proportion of life lived as of *T* collection date, and s() is the smoothing parameter. Each mouse had multiple data points across T collection date. This clustering was accounted for with a random intercept for ID, specified as (1|ID) above. We performed hypothesis tests related to the GAMM fits to explore trait sensitivity to bodyweight (m1 v. m2), PLL, (m1 v. m3), diet (m1 v. m4), and diet-by-trait interaction (m1 v. m5). Using the models specified above and a conservative false discovery rate (FDR < 0.01, method Benjamini-Hochberg), we thus identified traits that responded additively to body weight, traits that responded additively to diet, traits that responded additively to PLL (scaled age), and traits that responded interactively to diet and age. Traits included weekly body weights, biannual frailty, grip strength, and body temperature, and annual metabolic cage, body composition, echocardiogram, wheel running, rotarod, acoustic startle, bladder function, fasting glucose, immune cell profiling, and whole blood analysis (up to 689 measurements per mouse). Traits were categorized as related to health, metabolism, hematology, immune, or microbiome. For each trait category, barplots were generated to show the number of traits with significant (FDR < 0.01) associations with bodyweight (BW), diet, biological age (PLL), and diet x PLL interactions.

#### Lifespan prediction

To identify predictors of lifespan, we analyzed the correlation of 689 single-timepoint traits with lifespan after adjusting for effects of diet and bodyweight. We used a linear model: Lifespan ∼ Diet + BWTest + Trait. For body composition and change-in-bodyweight traits, we dropped the BWTest term. All continuous variables except PLL were rank normal scores transformed prior to model fitting.

#### Network modeling

A multivariate network analysis revealed which traits potentially mediate the effects of DR on lifespan. An empirical covariance matrix was estimated using the nonparanormal SKEPTIC estimator derived from the pairwise Kendall-Tau correlation using pairwise-complete data. The estimated covariance matrix was projected to the nearest positive definite matrix by truncating eigenvectors with negative eigenvalues. A sparse low-rank Gaussian graphical model was fitted to this covariance estimate using the ggLASSO python package with parameters lambda = 0.1, mu = 100. The result was normalized to obtain an inferred partial correlation matrix which we use as our phenotype network for downstream analysis.

To cluster the phenotypes, we constructed a weighted k-nearest neighbors (KNN) graph using the absolute partial correlation from above as similarity weights (i.e., we kept the k-largest weights per node) using k=10. The resulting KNN was clustered using spectral clustering to obtain 20 clusters which were labeled by hand. Diet and lifespan were broken out into their own univariate clusters and the body weight cluster was split into lean tissue mass and adiposity (fat mass / total mass) to account for body composition effects known from literature. This resulted in a summarized representation of 23 clusters (diet, lifespan, and 21 groups of traits).

We obtained a covariance decomposition by recomputing a sparse low-rank network (as above) on the reduced representation of 23 features. To obtain the relative importance of different effects of DR on lifespan, we performed covariance path decomposition^80^ of the covariance between diet and lifespan using this reduced representation graphical model. The absolute values of path scores were used to rank the relative importance of each path and normalized to sum to 1 to estimate the fraction of the covariance between DR and lifespan explained. To obtain network visualizations, the position of nodes and orientation of edges were determined by computing max-flow through the path network. We define the path network as the graph formed by taking the partial correlation network but reweighting edges to be the sum of all (absolute) path scores of paths that contain that edge.

#### QTL mapping

Genetic mapping analysis was carried out using the R package qtl2^81^. Genome scans for lifespan and RDW were carried out with diet and bodyweight at 6 months as additive covariates. Founder haplotype effects were estimated treating 8-state genotypes as random effects.

## Data Availability

All processed data, data analysis scripts, and supplementary data files have been deposited with FigShare (https://doi.org/10.6084/m9.figshare.24600255.v1).

## Acknowledgements

We thank the JAX Nathan Shock Center Animal and Phenotyping Core team for their assistance with animal husbandry, data collection, and curation. We thank the staff at Rockstep Inc., for their support with customization of our LIMS system, and Graham Ruby, Calico, for helpful discussions and suggestions. This project utilized the Jackson Laboratory Scientific Services.

## Author Contributions

Andrea Di Francesco: formal analysis, writing, project administration. Andrew G. Deighan: formal analysis, data curation. Lev Litichevskiy: formal analysis, writing. Zhenghao Chen: formal analysis, methodology. Alison Luciano: formal analysis, methodology. Laura Robinson: supervision, project administration. Gaven Garland: data collection, data curation. Hannah Donato: data collection, data curation. Will Schott: data collection, methodology. Kevin M. Wright: formal analysis, methodology. Anil Raj: formal analysis, methodology. G.V. Prateek: formal analysis, methodology. Martin Mullis: formal analysis, methodology. Warren Hill: formal analysis, methodology. Mark Zeidel: supervision, methodology. Luanne Peters: investigation, writing. Fiona Harding: formal analysis, methodology, writing. David Botstein: conceptualization, funding acquisition. Ron Korstanje: supervision, project administration. Christoph A. Thaiss: supervision, writing. Adam Freund: conceptualization, supervision, project administration. Gary A. Churchill: conceptualization, funding acquisition, project administration, formal analysis, writing.

